# Mycobacterial surface shedding drives bystander cells response during early intracellular infection

**DOI:** 10.1101/2025.11.13.687818

**Authors:** Davide D’Amico, Lyudmil Raykov, Thierry Soldati, Mélanie Foulon

**Affiliations:** Département de Biochimie, Faculté des Sciences, Université de Genève, Sciences II, 30 quai Ernest Ansermet, CH-1211 Genève, Switzerland

**Author notes:** Co-corresponding authors &.

**Keywords:** *Mycobacterium marinum*, *Dictyostelium discoideum*, phagocytosis, cell-autonomous defense, envelope

## Abstract

Mycobacteria possess a complex cell envelope whose outermost layer plays an underexplored role in host-pathogen interactions. Using *Dictyostelium discoideum* as a model host phagocyte, we show that *Mycobacterium marinum* rapidly sheds its envelope components, including surface proteins, carbohydrates, and virulence-associated lipids, within minutes of uptake. Shed material is actively trafficking within host endocytic pathways and disseminates to neighboring bystander cells, where it accumulates and triggers several responses. Notably, bystander cells exposed to shed material exhibit a transient delay in G1/S cell cycle progression, an activation of membrane damage-response pathways, and an enhanced resistance to subsequent mycobacterial infection. These phenotypes are recapitulated by infection-free conditioning with purified envelope extracts, demonstrating that superficial components of the envelope alone are sufficient to modulate host cell responses. Moreover, this priming effect is independent of bacterial viability or the Esx-1 secretion systems, underscoring the intrinsic immunomodulatory capacity of the envelope. Interestingly, bacteria that lose their outer layer are more frequently ubiquitinated, suggesting that host-driven stripping exposes molecules that are recognized by cytosolic sensors to mount a cell-autonomous defense. Together, our findings reveal that mycobacterial envelope shedding is a widespread, early event during intracellular infection that impacts both infected and bystander cells. These findings suggest that mycobacterial outermost envelope components can influence host cell physiology and contribute to early innate immune modulation, with implications for understanding the initial determinants of infection outcomes.

## Introduction

Mycobacteria are a highly diverse group of bacteria that have evolved from environmental organisms to strict human pathogens through reductive evolution (Han & Silva, 2014). The most well-known pathogenic species for human include *Mycobacterium tuberculosis*, the causative agent of tuberculosis, *M. leprae*, which causes leprosy or *M. ulcerans*, responsible for Buruli ulcer. While these pathogens are characterized by their extremely slow growth, the rising emergence of non-tuberculous mycobacteria (NTM) infections, caused by faster-growing species such as *M. abscessus*, demonstrates that mycobacterial pathogenicity is not only linked to a slow metabolism. One of the distinctive features of *Mycobacterium* species is their unique cell envelope, which differs from that of Gram-positive or Gram-negative bacteria. This complex protection contributes to the bacteria’s ability to survive in hostile environments (Daffé & Marrakchi, 2019). The envelope is composed of several layers, including peptidoglycan, covalently linked to arabinogalactan, which is esterified by long-chain mycolic acids. Mycolic acids form the inner leaflet of an outer membrane bilayer, contributing to the bacterial cell’s impermeability and rigidity. In addition, mycobacteria possess an outermost layer, often referred to as the “capsule”. This layer, primarily thought to be a specific feature of pathogenic *Mycobacterium* species (Daffé & Etienne, 1999; Fréhel, 1988), has been identified in a broader spectrum of pathogenic and non-pathogenic, slow or fast-growing mycobacteria (Sani et al., 2010). This labile layer, composed of a mixture of glycans, lipids and proteins, can vary in thickness, reaching up to 40 nm in *M. marinum* (Sani et al., 2010). Despite the recognition of its diversity in biogenesis and composition, the role of the outermost cell envelope layer in mycobacterial interactions with the host remains poorly understood.

Over the past two decades, a growing body of evidence has revealed that mycobacteria release and shed their cell wall components during host infection, contributing to both immune modulation and pathogenesis. Beatty et al. (Beatty et al., 2000; Beatty et al., 2001) were first to demonstrate that fluorescently labeled cell wall surface components (glycoconjugates and proteins) traffic from the phagosome into host endocytic compartments and are ultimately released through exocytosis. Complementary findings showed more specifically that mycobacterial lipids such as lysocardiolipin (Fischer et al., 2001) and lipoproteins, such as the 19-kDa Ag lipoprotein (Neyrolles et al., 2001), dissociate from intra-phagosomal mycobacteria and follow independent trafficking routes within macrophages. While all these studies were performed with the attenuated *M. bovis* BCG strain, it was later shown that also virulent intracellular mycobacteria, such as *M. marinum*, can shed cell wall fragments after escape from the phagosome to the cytosol (Carlsson et al., 2009). These foundational insights were later extended by studies showing that mycobacteria release not only structural components but also nucleic acids, such as DNA sensed by the cytosolic DNA sensor cyclic GMP-AMP synthase (cGAS) (Manzanillo et al., 2012; Watson et al., 2015), and that certain cell wall lipids like PDIM can transfer between infected and uninfected host cells (Cambier et al., 2020). Together, these findings sporadically outline an interplay between the pathogen and the host beyond the confines of the mycobacteria-containing vacuole. But the dynamic and mechanistic aspects of envelope components shedding remained to be elucidated.

In this context, we used *Dictyostelium discoideum* as a simple cellular model to investigate the early stages of mycobacteria phagocytosis. As a professional phagocyte, it shares similarities with mammalian innate immune cells in both morphology and behavior, making it a suitable tool for investigating host-pathogen interactions, including mycobacterial infections (Cardenal-Muñoz et al., 2018; Dunn et al., 2017; Guallar-Garrido & Soldati, 2024). In recent years, *D. discoideum* has emerged as a powerful and experimentally versatile model host organism to decipher complex human diseases like Alzheimer’s or Parkinson’s (Chen et al., 2017; Frej et al., 2017; Gilsbach et al., 2012; Ludtmann et al., 2014), to investigate infections by several pathogens, such as *Legionella pneumophila*, *Mycobacterium spp*., *Vibrio cholera*, *Francisella noatunensis*, *Pseudomonas aeruginosa*, *Salmonella enterica*, yeasts, and fungi like *Cryptococcus neoformans* and *Aspergillus fumigatus* (reviewed in Cardenal-Muñoz et al., 2018; Dunn et al., 2017), as well as to perform high-throughput genetic and drug screens (Hanna et al., 2020; Liao et al., 2016; Nitschke et al., 2024; Ouertatani-Sakouhi et al., 2017).

In this study, we show that *M. marinum* outermost layer is stripped from its surface upon internalization in *D. discoideum*, filling the phagosomal lumen. We observed that the superficial components of the *M. marinum* envelope, including surface-exposed carbohydrates, proteins, and virulence-associated factors are rapidly released within the first 4 hours post-infection. This passive shedding is conserved across different mycobacterial species and occurs concurrently with significant remodeling of the MCV. Furthermore, we demonstrate that shed material actively traffics within the infected cell’s endosomal compartment in an ESCRT-dependent manner and is disseminated to bystander cells, where it delays the cell cycle, particularly at the G1 to S transition, and primes them to better resist subsequent infection. These findings highlight the dynamic role of cell wall shedding in host-pathogen interactions and indicate that mycobacterial envelope material can influence the host cell autonomous immune response by affecting both infected and bystander cells.

## Results

### The superficial layer of the *M. marinum* cell wall is shed early during phagocytosis

To investigate the dynamics of *M. marinum* cell wall during infection, surface-exposed carbohydrates and proteins were respectively labeled with AlexaFluor-hydrazide (after Schmidt reaction to form reactive aldehyde residues) or AlexaFluor-NHS (by ester bioconjugation to amine groups), following procedures established in (Beatty et al., 2000) (**Figure 1A**). Labeling efficiencies were monitored by flow-cytometry (**Suppl. Figure 1A-B**), and the viability of labeled bacilli was verified by counting colony-forming units (CFU) (**Suppl. Figure 1C**). Early events of phagocytosis were observed by high-resolution microscopy after spinoculating pre-labeled bacteria onto *D. discoideum* cells. As shown in **Figure 1B**, 10 min post-contact between cells and bacteria, AlexaFluor-488 NHS already fills the bacilli-containing compartment, indicating a fast release of bacterial surface components into the phagosomal environment. Similar observations were made with BV-2 microglial cells (another phagocyte model), for which the very first step of bacilli entry was captured (**Figure 1C-D**). In those cells, in the first 40 min of phagocytosis, *M. marinum* surface-labeled components appeared to be stripped and trafficked inside the host cell. When observed slightly later during the infection in *D. discoideum* (2 hours post-infection (hpi), early phase of the MCV genesis), both surface-labeled carbohydrates and proteins were found accumulating in enclosed compartments at some distance from the MCV (**Figure 1E**). Immunofluorescence staining of fixed infected cells at 2 hpi highlighted a co-localization of AlexaFluor-488 hydrazide labeled shed material with an anti-PGL antibody (**Figure 2A**). Phenolic GlycoLipid (PGL) is a mycobacterial lipid known to play a role in the virulence of several pathogenic mycobacteria, including *M. leprae* and *M. tuberculosis*. To further confirm its shedding from *M. marinum* cell wall during phagocytosis, we performed a metabolic pre-labeling of the bacteria (according to (Guzmán et al., 2024)) and infected *D. discoideum* cells as above. PGL labeling was verified by thin-layer chromatography (TLC), to exclude any aspecific labeling of closely related lipid species, such as PDIM (**Suppl. Figure 1D-E-F**). As for surface-exposed carbohydrates or proteins, prelabeled-PGL was released from *M. marinum* surface and accumulated in small intracellular compartments away from the MCV (**Figure 2B**). Altogether, these observations show that shed envelope material, containing virulence-associated factors, diffuses away from the bacteria and is transported from the MCV in the earliest phase of phagocytosis.

**Figure 1.**
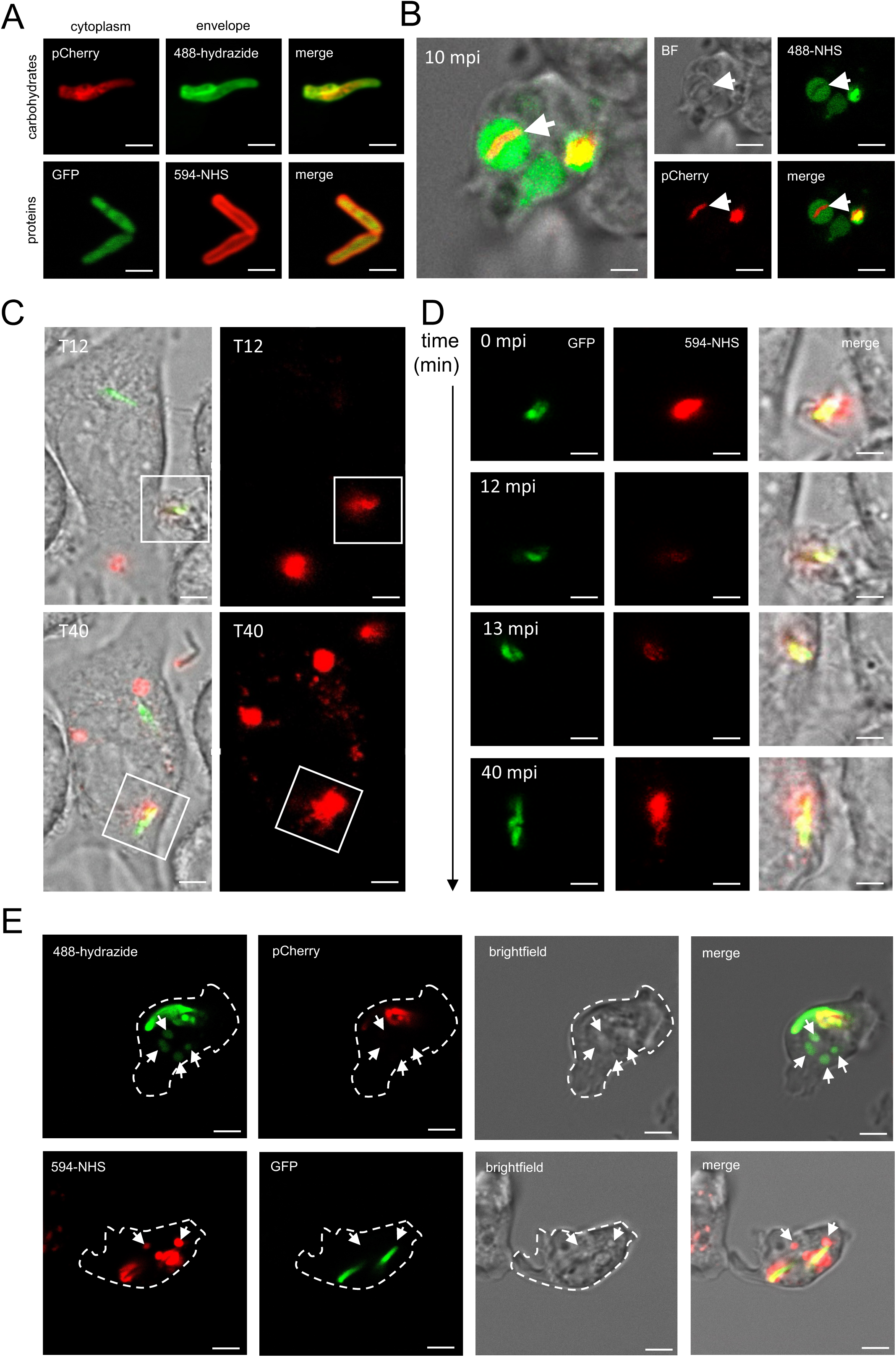
*M. marinum* surface-exposed carbohydrates and proteins are shed during phagocytes infection. (**A**) High-resolution images of *M. marinum* expressing cytoplasmic pCherry or GFP respectively labeled with 488-hydrazide or proteins with 594-NHS shows surface-specific labeling. Scale bars = 0.5 µm. (**B**) High-resolution live acquisition of *D. discoideum* infected cells 10 min post-infection, highlights *M. marinum* (expressing cytoplasmic pCherry) labeled surface components (488-NHS) shedding and filling the bacteria-containing compartment (white arrow). Scale bars = 2 µm. (**C**) Live acquisition of BV-2 cells phagocyting *M. marinum* GFP labeled for surface-exposed proteins (594-NHS), showing the entry (from 0 to 13 minutes post-infection, mpi) and stripping (at 40 mpi) of *M. marinum* labeled surface components during the first 40 min (T40) of phagocytosis. Scale bars = 5 µm for enlarged view of the cell, and 1 µm for zoomed panels. (**D**) High-resolution live acquisition of *D. discoideum* infected cells at 2hpi, showing shed material (488-hydrazide or 594-NHS labeled) accumulating in some cellular compartments (white arrows) at distance from the bacilli (seen with pCherry or GFP cytoplasmic reporters). Scale bars = 5 µm.

**Figure 2.**
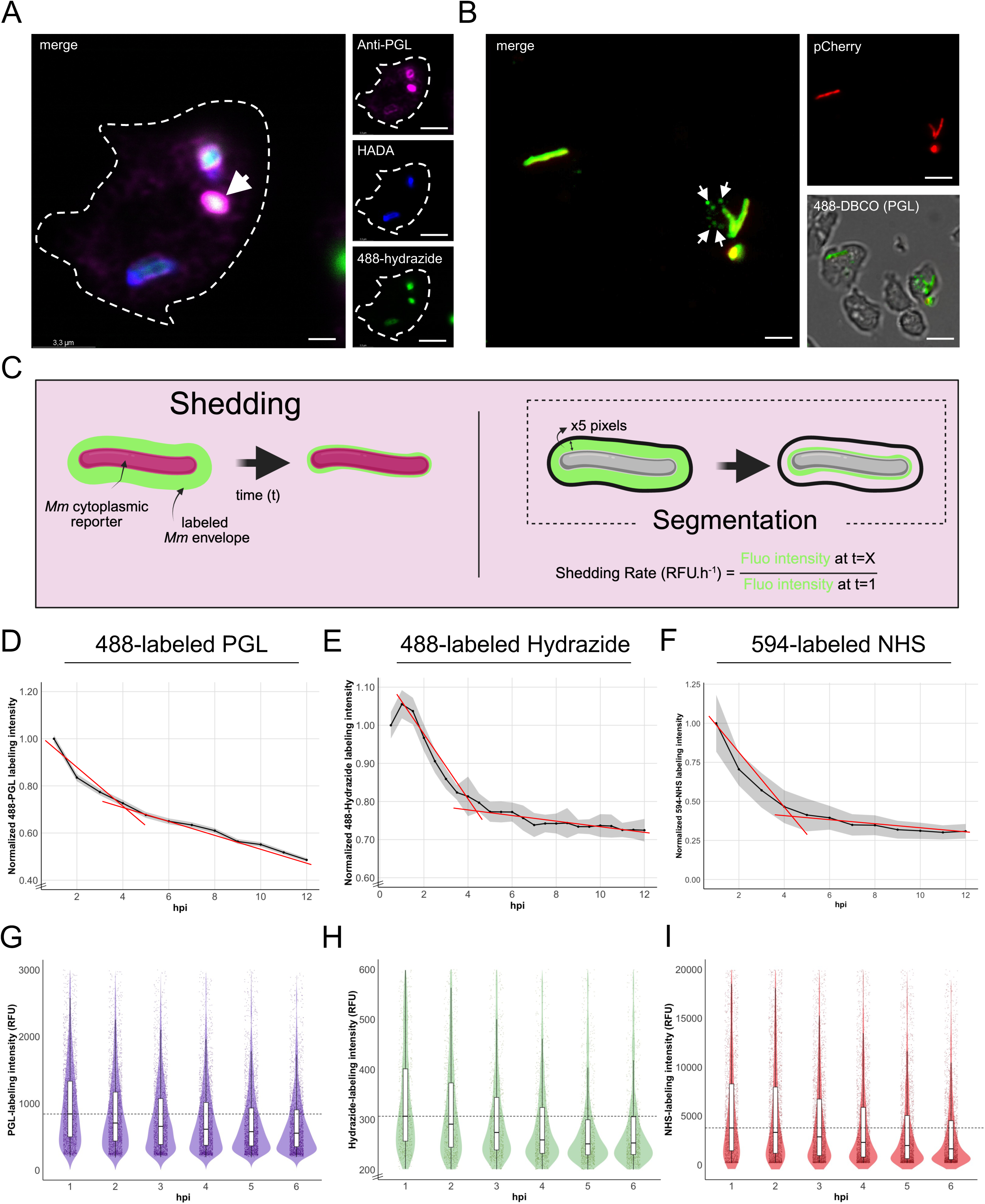
*M. marinum* shed material, containing the virulence associated factor PGL, is rapidly released during the establishment of the MCV. (**A**) Immunofluorescence staining on fixed *D. discoideum* infected cells at 2hpi shows a co-localization of 488-hydrazide labeled shed material with an anti-PGL antibody (white arrow). Scale bars = 1.5 µm for enlarged merged view and 5 µm for individual channel images. (**B**) Live acquisition of *D. discoideum* cells infected with *M. marinum* pCherry metabolically pre-labeled for PGL shows 488-DBCO-clicked PGL shedding (white arrows) at 2hpi. Scale bars = 2.5 µm for enlarged merged view and 10 µm for individual channel images. (**C**) Schematic representation of shedding parameters acquired and quantified by high-content microscopy. The fluorescence intensity of the labeled envelope was measured in the extended bacterial mask. Quantification by high-content microscopy shows a two-phases shedding dynamic (fast until 4hpi, slower until 12hpi) for (**D,G**) 488-DBCO labeled PGL, (**E,H**) 488-hydrazide labeled surface exposed carbohydrates and (**F,I**) 594-NHS labeled surface exposed proteins. Ratio to the median of initial fluorescence intensity in the bacterial mask are showed in panels **D**, **E** and **F**, while raw fluorescence intensities (expressed as relative fluorescent units (RFU)) for each segmented intracellular bacterial event is depicted as violin plots in panels **G**, **H** and **I**. Data in panels **D**, **E** and **F** are represented as median +/- SEM, with linear regression models as red lines. Dashed line in panels **G**, **H** and **I** shows median for the first time-point (1hpi). Results are shown as representative of two to three independent replicates.

### *M. marinum* cell wall superficial layer shedding happens in the first 6hpi

The dynamic of *M. marinum* cell wall superficial components shedding during the infection in *D. discoideum* was monitored by high-content microscopy over 12 hpi, and shedding rates were quantified accordingly (**Figure 2C**). During this time-lapse, *M. marinum* is not replicating yet as reflected by CFU counting or RFU measurement (Hagedorn & Soldati, 2007; **Suppl. Figure 1G**), ensuring no dilution of the labeled signal due to bacterial division. Cells were infected with *M. marinum* labeled for PGL (**Figure 2D-G**), surface-exposed carbohydrates (**Figure 2E-H**) or proteins (**Figure 2F-I**) and the intensity of fluorescence (corresponding to the labeled components) was measured in the extended bacterial mask (**Figure 2C**). Shedding kinetics, reflected by the loss of fluorescence intensity in the mask, was overall similar for the three types of labeling, with a first phase of drastic decrease from 1 to 4 hpi followed by a second phase showing a milder decrease up to 12 hpi (**Figure 2D-E-F**). During this first phase, a loss in intensity of approximately 35% was observed for PGL (**Figure 2D**), with a normalized rate of 0.08 RFU.h^-1^.Similarly, a loss of intensity of around 22% (**Figure 2E**), with 0.06 RFU.h^-1^ was quantified for surface-exposed carbohydrates. The release of surface-exposed proteins occurred the fastest, with a loss of almost 70% (**Figure 2G**), with a normalized rate of 0.18 RFU.h^-1^ up to 4 hpi (**Figure 2F**). A certain heterogeneity was observed for the three types of pre-labeling at 1 hpi. This heterogeneity tended to diminish by 6 hpi (**Figure 2G-H-I**). These results show that *M. marinum* sheds its cell wall rapidly within the first 4-6 hpi. This early phase coincides with significant remodeling of the phagosomal compartment (Cardenal-Muñoz et al., 2018; **Suppl. Figure 1H**). Therefore, cell envelope superficial shedding may play a crucial role in the formation of the MCV, which is essential for successful infection with *M. marinum*.

### Cell wall shedding is conserved among mycobacteria species, but the dynamics depends on mycobacterial intracellular compartmentalization

We showed that the shedding of superficial components of mycobacterial cell wall is conserved among two evolutionary distant phagocyte models (**Figure 1**). We then asked whether this phenomenon was also conserved between mycobacteria species. Several pathogenic (*M. kansasii*, *M. abscessus* and *M. tuberculosis*) and attenuated or non-pathogenic strains (*M. bovis* BCG, *M. smegmatis* and *M. brumae*) were assessed. Their peptidoglycan was metabolically pre-labeled with the fluorescent precursor NADA (3-[7-nitrobenzofurazan]-carboxamide-D-Alanine) and surface-exposed proteins were labeled with AlexaFluor-594 NHS following the procedure used for *M. marinum* labeling. Those pre-labeled mycobacteria were used to infect *D. discoideum*, and infected cells were observed by high-resolution microscopy at 2 hpi (**Figure 3A to C**). Shedding of labeled surface-exposed proteins was observed for all species, pathogenic or not, fast or slow growers. This shedding was specific for the outermost cell wall layer because no release of NADA-prelabeled peptidoglycan was observed for any *Mycobacterium* species. We then assessed whether shedding dynamics could be affected by mycobacterial virulence. To that end, shedding rates were measured by high-content microscopy during the infection, comparing *M. marinum* WT and ΔRD1. *M. marinum* ΔRD1 (Region of Difference 1) is an attenuated mutant that does not assemble its Esx-1 system involved in phagosomal damage via the secretion of the membrane-damaging toxin EsxA. Damage is essential for *M. marinum* to efficiently establish its MCV and later escape to the cytosol. Interestingly, while the first minutes after uptake were similar (**Suppl. Figure 2F**), *M. marinum* ΔRD1 sheds its superficial prelabeled proteins and carbohydrates faster than WT, as reflected by higher shedding rates in the first phase (**Figure 3D to F and 3G to I**). Importantly, pre-labeling efficiency was identical for both strains and both types of labeling (**Suppl. Figure 2A-C** for surface-exposed carbohydrate labeling, i.e. AlexaFluor-488 hydrazide, and **Suppl. Figure 2B-D** for surface-exposed proteins labeling, i.e. AlexaFluor-594 NHS) as well as their viability (**Suppl. Figure 2E**). More specifically, bacteria unable to inflict membrane damage occupy MCV that remain efficiently connected to endosomal membrane trafficking (Kreibich et *al.* 2015) favoring the release and trafficking of their superficial layer. To test this hypothesis, we therefore similarly monitored shedding rates during *M. marinum* WT infection in *D. discoideum* Δatg1, fully impaired in autophagy, or Δalix, affected in ESCRT-mediated mechanisms. These cells are unable to effectively repair the damage induced by *M. marinum* to the MCV membrane, leading to an early escape of the bacilli to the cytosol (Guallar-Garrido & Soldati, 2024). For both surface-exposed proteins (**Figure 3J to L**) or carbohydrates (**Figure 3M to O**), shedding rates were slower in *D. discoideum* Δatg1 and Δalix cells, with the most drastic effect being observed for surface-exposed proteins in Δalix cells with a shedding rate of 0.04 RFU.h^-1^ against 0.23 RFU.h^-1^ in WT cells. Altogether, the results indicate that intracellular compartmentalization and endolysosomal membrane trafficking drive the release dynamics of *M. marinum* surface components.

**Figure 3.**
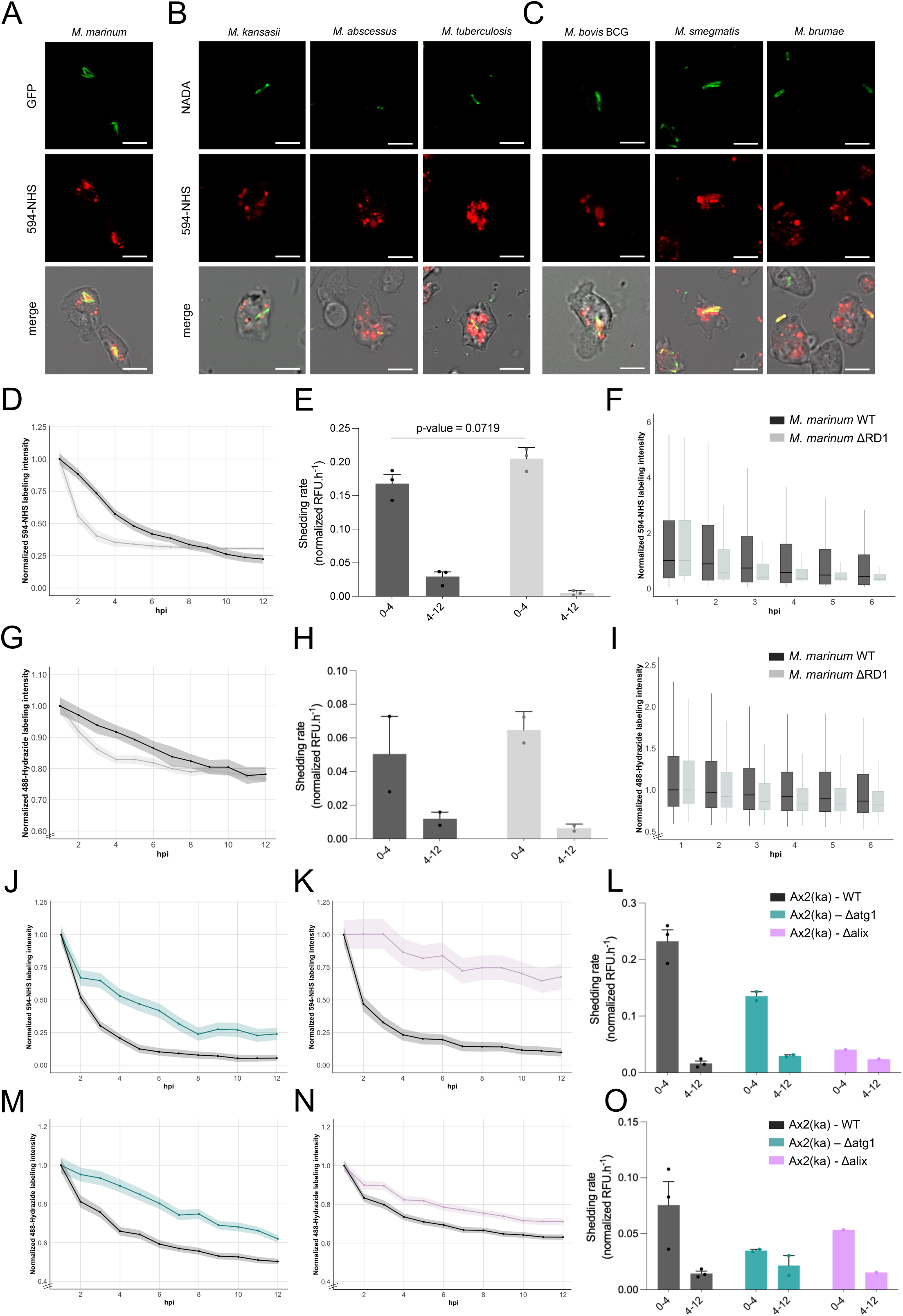
Mycobacterial cell wall superficial layer shedding is conserved, but the dynamics depends on virulence. Shedding of 594-NHS labeled surface components is observed during infection of *D. discoideum* cells with several pathogenic (**A,B**) (*M. marinum*, *M. kansasii*, *M. abscessus* and *M. tuberculosis*) or (**C**) non-pathogenic (*M. bovis* BCG, *M. smegmatis* and *M. brumae*) mycobacteria. Peptidoglycan, metabolically pre-labeled with NADA, shows no shedding of deeper cell wall layer at 2hpi. Scale bars = 5 µm. Quantification by high-content microscopy of (**D**) 594-NHS or (**G**) 488-Hydrazide labeling intensity shows a faster shedding of surface-exposed proteins and carbohydrates, respectively, for *M. marinum* ΔRD1 (grey) when compared to *M. marinum* WT (black) during the first 6hpi in *D. discoideum.* (**E,H**) Shedding rates were estimated based on the slopes of the curves from 0 to 4 and 4 to 12 hpi for each condition. (**F,I**) The faster decrease in normalized fluorescence intensity is reflected at population level for both conditions. Quantification by high-content microscopy of (**J,K**) 594-NHS or (**M,N**) 488-Hydrazide labeling intensity shows a slower shedding of surface-exposed proteins and carbohydrates, respectively, in *D. discoideum* cells altered in autophagy response (Δatg1) or in escrt-mediated responses (Δalix). (**L,O**) Shedding rates were estimated based on the slopes of the curves from 0 to 4 and 4 to 12 hpi for each condition. Data in panels **D, G, J, K, M** and **N** are represented as median +/- SEM. Data in panels **F** and **I** are depicted with box-plots with median and +/- SD. Results are shown as representative of at least three independent experiments. Data in panels **E, H, L** and **O** are represented as bar plots + SD, with individual dots representing three independent biological replicates. Results are shown as representative of at least three independent experiments.

### Shed mycobacterial material traffics in endosomal compartments

In mammalian macrophages, mycobacterial cell wall components were shown to traffick throughout the endocytic compartment (Beatty et al., 2001; **Suppl. Figure 3C-D**). In *D. discoideum* phagocytes, *M. marinum* labeled carbohydrates and proteins were observed in enclosed compartments budding from the MCV at 2 hpi (**Figure 4A-B**). Immunofluorescence staining on fixed cells showed that this shed material can be found in p80-positive compartments (**Figure 4C**), a ubiquitous marker of endosomes and lysosomes, confirming the observation above. Moreover, the use of an anti-ubiquitin antibody (FK2) showed that while shed material was not significantly positive for ubiquitin, bacteria poorly or not anymore labeled for their surface-exposed proteins tended to be highly ubiquitinated (**Figure 4D**). In contrast, *M. marinum* still covered with their pre-labeled superficial layer were not ubiquitinated (**Figure 4D**). This might reflect either bacteria still confined in their MCV, or cytosolic bacteria having lost their surface-exposed proteins normally sensed and ubiquitinated. Together, these observations support an active dissemination of the mycobacterial superficial layer driven by host’s membrane trafficking. This removal might allow a better access to components of the cell wall recognizable by the host defense machinery.

**Figure 4.**
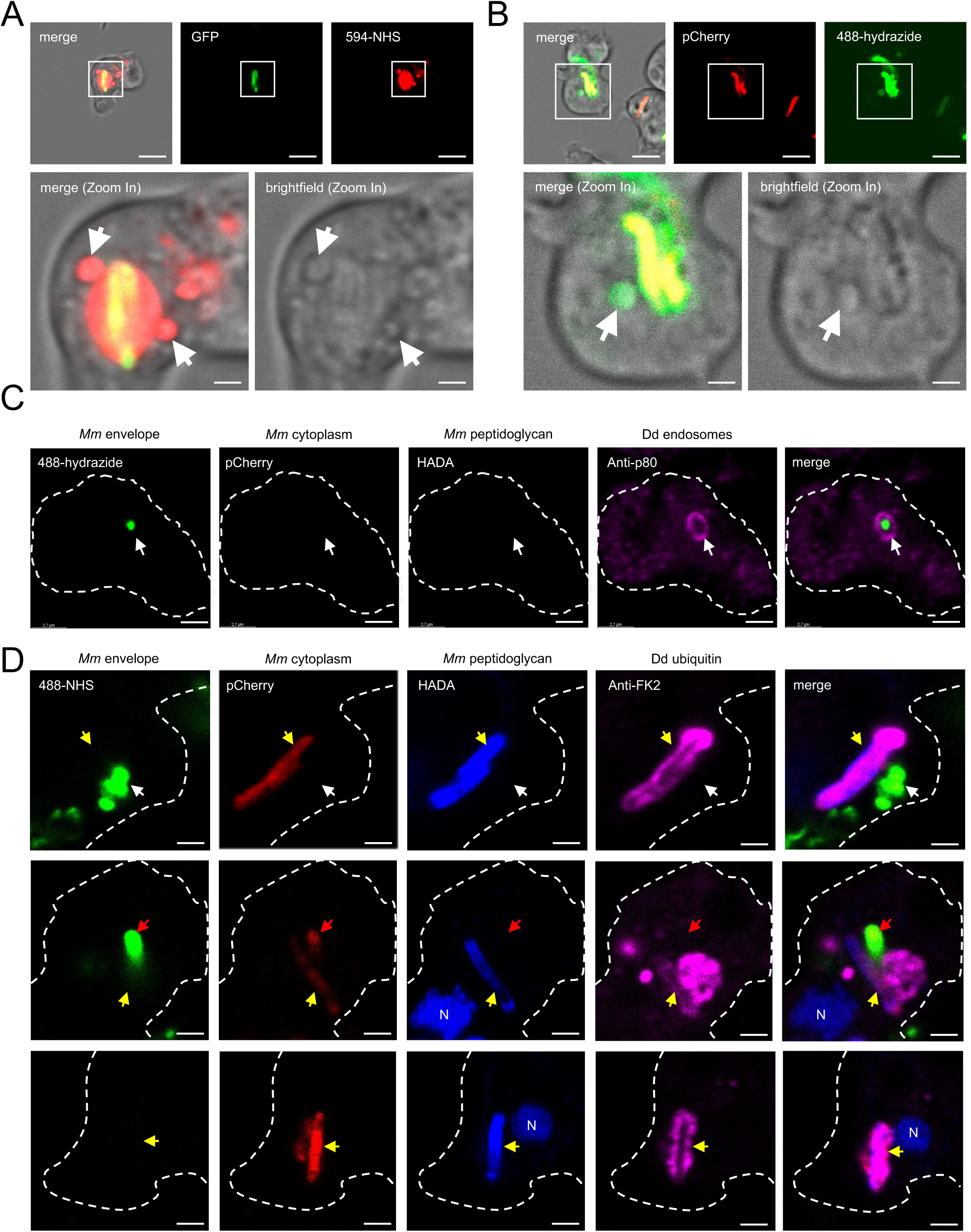
*M. marinum* shed material is trafficking in endosomal compartments, while stripped bacteria are more ubiquitinylated. High-resolution images captured at 2hpi in *D. discoideum* cells infected with *M. marinum* WT pre-labeled with 594-NHS (**A**) or 488-hydrazide (**B**) show accumulation of labeled material in enclosed compartments visible on the brightfield (white arrows). Scale bars = 8 µm, and 0.5 µm in inset images, for panel **A**, and 5 µm and 0.5 µm in inset images for panel **B**. (**C**) Immunofluorescence staining on fixed *D. discoideum* cells infected with *M. marinum* WT expressing a cytoplasmic pCherry reporter, pre-labeled for surface-exposed carbohydrates (488-hydrazide) and peptidoglycan (HADA) shows the presence of shed material in p80-positive endosomal compartment (white arrow) at 2hpi. Scale bars are 2 µm. (**D**) Immunofluorescence staining with an anti-ubiquitin (FK2) antibody shows no ubiquitinylation of shed labeled proteins (488-NHS) (white arrow) but intense ubiquitinylation of stripped *M. marinum* (bacilli having already lost their labeled superficial layer) (yellow arrows). *M. marinum* bacilli still labeled with 488-NHS are not recognized by the anti-ubiquitin (red arrow). Mm = *M. marinum*, Dd = *D. discoideum*, N = Nuclei. Scale bars = 0.5 µm.

### The shed material is actively disseminated to bystander cells

During live-microscopy recording of *D. discoideum* cells infected with AlexaFluor-594-NHS-labeled *M. marinum*, transfer of envelope material was observed from infected to non-infected cells (**Figure 5A**). This observation suggested that this material disseminates to bystander cells. To test this hypothesis quantitatively, we designed a dissemination assay (**Figure 5B**), in which an infected *D. discoideum* population (donor) expressing cytosolic mCherry (act5::mCherry) was mixed with an unmarked non-infected population (acceptor) at a 1:1 ratio. In this assay, AlexaFluor-488-NHS-labeled *M. marinum* also expressing cytosolic mCherry was used to infect the donor population. Accumulation of AlexaFluor-488-NHS in acceptor cells was measured by high-content microscopy over 6 hours post-contact (**Figure 5C**). The proportion of 488-NHS positive acceptor cells (reflected by a signal intensity at 510 nm above baseline, therefore named as proportion of positive-cells) increased over time, reaching 8% 6 hours after contact (**Figure 5D-E**), indicating net transfer of shed material without associated mCherry bacteria to non-infected cells early after contact. Another observation was made during *D. discoideum* infection with pre-labeled *M. marinum*. When looking at large fields of view, some bystanders not only received but seemed to accumulate 594-NHS labeled shed material (**Figure 6A**). To quantify the trafficking and dissemination of shed material within the host cells, the fluorescence intensity, reflecting labeled *M. marinum* shed material, was measured in the cell mask excluding the extended bacterial mask (**Figure 6B**), and normalized to the median fluorescence intensity in mock-treated cells (**Figure 6C to F**). Quantification of the accumulation of shed material was applied to the two subsets of the population: bystander and infected cells from *D. discoideum* WT (**Figure 6C**), Δatg1 (**Figure 6D**) or Δalix (**Figure 6E**) cells infected with *M. marinum* WT. For cells infected with *M. marinum* ΔRD1, mock-treated conditions were missing, therefore normalization was applied as the proportion of cells above baseline values (**Figure 6F**). Trafficking rates (i.e. fluorescence intensity in the cell mask excluding bacteria extended mask, over the time) were measured for all the conditions and cell subsets (**Figure 6G**). A fast decrease in fluorescence from 1 to 3 hpi was observed in WT cells infected with *M. marinum* WT or ΔRD1, as well as for Δatg1 cells, happening concomitantly with an increase of the signal in bystander cells. In contrast, little change was observed over the time-course in Δalix cells as reflected by reduced trafficking rates in those cells when compared to WT cells (**Figures 6H-I**). Dissemination ratio, reflecting the transfer of shed material from infected to bystander cells (**Figure 6J**) were compared as well. All ratio were positive except for Δalix cells, showing a defect in transfer of *M. marinum* shed material from these cells. It is to note that similar quantifications with 488-hydrazide labeled *M. marinum* surface components slightly differed, with defective trafficking but similar dissemination in both Δatg1 and Δalix when compared to WT cells (**Suppl. Figure 4**). While interesting, this difference might come from a technical bias due to the low signal intensity of 488-hydrazide labeling and the high background in the green channel in *D. discoideum* cells. Overall, this shows that *M. marinum* shed material, more notably proteins, which actively traffic, disseminate to and accumulate in bystanders early during infection.

**Figure 5.**
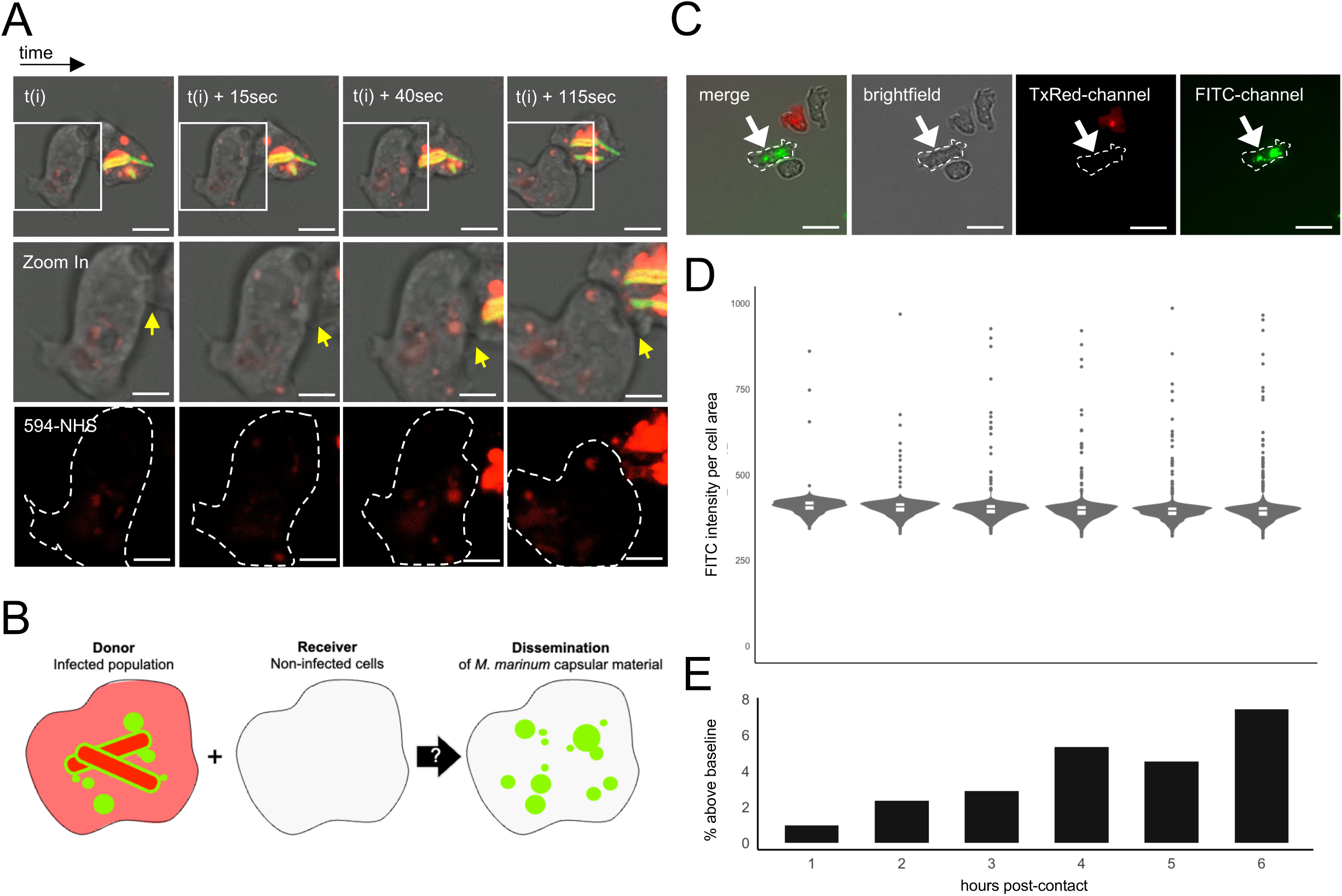
*M. marinum* shed material is actively disseminated. (**A**) Live microscopy recording of *D. discoideum* infected populations with *M. marinum* pre-labeled with 594-NHS at 2hpi shows transfer of shed material from an infected cell to its neighboring non-infected cell (yellow arrow), without bacteria cell-to-cell spreading. Scale bars = 8 µm, and 5 µm for zoomed in images. (**B**) Schematic representation showing dissemination assay experimental design. Briefly, *D. discoideum* cells expressing a cytosolic mCherry reporter were infected with *M. marinum* also expressing a cytoplasmic pCherry reporter and pre-labeled with 488-NHS. Infected population (donors) was mixed with non-infected *D. discoideum* cells (receivers) and presence of 488-NHS labeled material was quantified in this subset. (**C**) A receiver cell (dashed line with white arrow) shows accumulation of 488-NHS labeled shed material, without showing any bacterium (pCherry) inside. Scale bars = 5 µm. (**D**) FITC intensity (reflecting 488-NHS signal) measured per cell (RFU) increases in a small subset of the *D. discoideum* receiving cell population up to 6hours post-contact, as reflected by the percentages of cells with FITC value above baseline (**E**).

**Figure 6.**
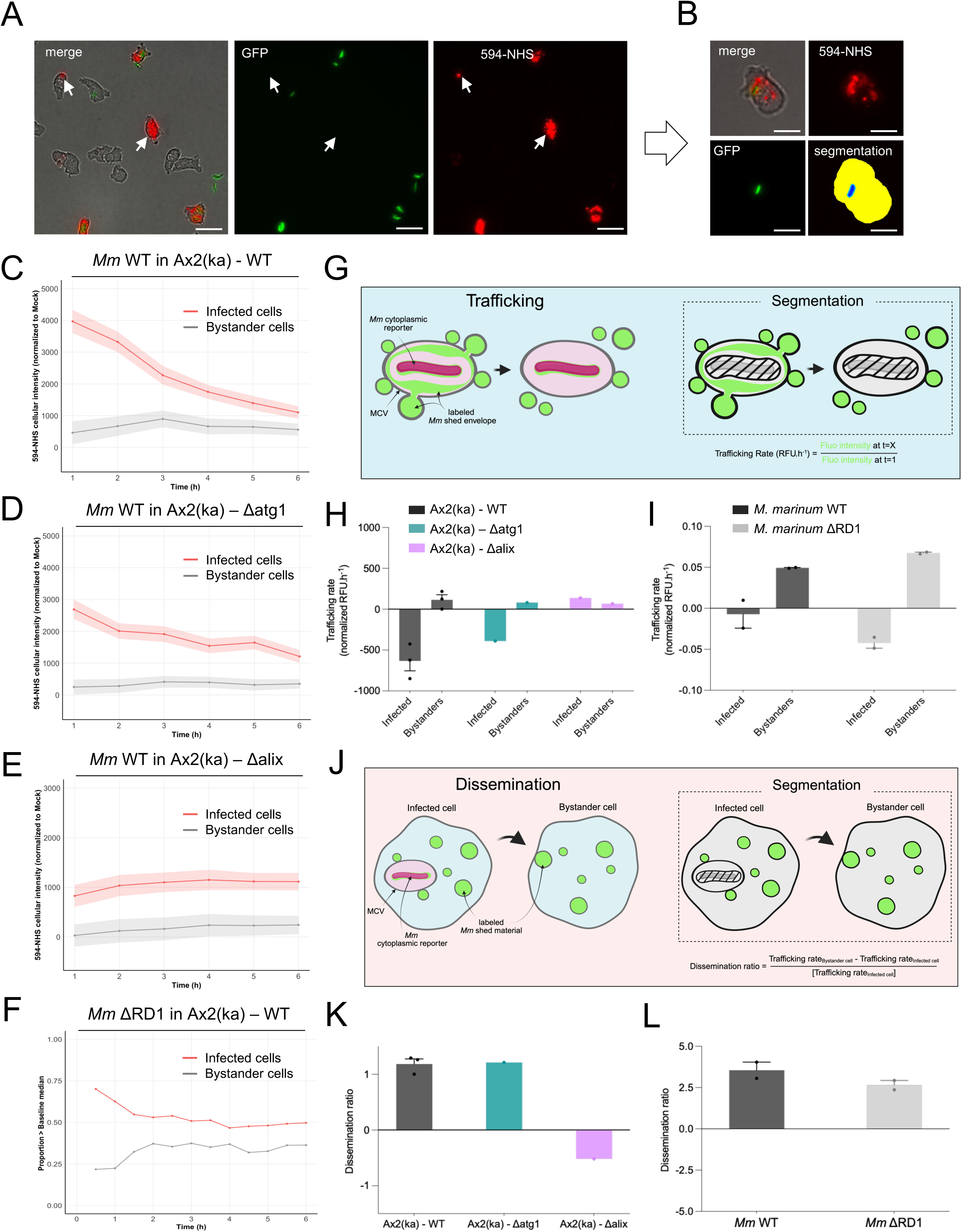
*M. marinum* shed material is actively trafficking and disseminating to bystander cells. **(A)** In *D. discoideum* infection, large field of view shows accumulation of 594-NHS labeled shed material in non-infected cells (white arrows) at 2hpi. Scale bars = 20 µm. (**B**) High-content microscopy-based segmentation allows to measure shed material accumulation in the cell cytosol (yellow surface) by excluding the extended bacterial mask (blue surfaces). Total fluorescence intensity labeled *Mm* shed material (594-NHS) was measured in infected (red lines) and bystanders (grey lines) cells, and normalized to the median in mock for *D. discoideum* WT (**C**), Δatg1 (**D**) or Δalix (**E**) cells infected with *M. marinum* WT. Normalization to max baseline was applied for *D. discoideum* WT cells infected with *M. marinum* ΔRD1, as mock conditions were missing in those infections (**F**). Therefore, proportions of events above the established baseline are depicted for this panel. Results (median +/- SEM) are shown as representative of one to three independent experiments. (**G**) Schematic representation of trafficking parameters quantified by high-content microscopy. The fluorescence intensity of the labeled shed material was measured in the cell mask, excluding the extended bacterial mask. (**H,I**) Trafficking rates in infected and bystander cells were calculated based on the first 3 hpi. **(J**) Schematic representation of dissemination parameters quantified by high-content microscopy. Dissemination ratios were measured based on trafficking rates previously calculated. (**K,L**) Dissemination ratios were calculated for each condition. Results are shown as bar plots +SD, with individual dots representing three independent biological replicates.

### The shed envelope material delays the cell cycle of bystander cells

To investigate the impact of this accumulation of shed material in bystander cells, a microfluidic device (Mottet et al., 2021) was used for long-term single-cell imaging of the two cellular subsets during infection. Briefly, after infection of *D. discoideum* with GFP-expressing *M. marinum*, cells were trapped in microchambers, and high-content microscopy was performed to monitor cell division (**Figure 7A**). Comparisons between mock-treated, bystander and infected cells highlighted an interdivision time prolonged approximatively two-fold in cells infected with *M. marinum* WT (17.18 h against 8.24 h for mock cells), and 1.8-fold when infected with *M. marinum* ΔRD1 (15.33 h). Interestingly, bystanders from both infections showed, as well, a delay in their cell cycle, with an interdivision time of 13.39 h and 13.03 h, respectively. More importantly, cells phagocytosing antibiotic-killed *M. marinum* also showed an interdivision time prolonged by 1.5-fold, showing that an active bacterial replication or metabolism is not required to induce cell cycle alterations in host cells. This correlates with previous observation of cell wall shedding from dead *M. bovis* BCG in macrophages (Rhoades et al., 2003). In *D. discoideum*, the cell cycle is divided in the standard M, G1, S and G2 phases with a very short G1 and a prolonged G2 that lasts almost 70% of the total (Zada-Hames & Ashworth, 1978; **Figure 7B**). To decipher which cell cycle check point is impacted by the infection, we performed RNA-sequencing in mock-treated and infected populations at 4 hpi and interrogated different lists of genes known to be associated to the transitions between each phase. Compared to the strong upregulation of genes in mock-treated cells that divide normally, a strong down-regulation of gene expression related to the G1 to S transition was observed in *M. marinum* WT infected populations, and, to a milder extent, in *M. marinum* ΔRD1 infected populations and cells in contact with dead bacilli (**Figure 7C**). This trend was also observed in genes related to the M to G1 transition (**Suppl. Figure 5A**). To better investigate these effects in the specific cell subsets, FACS-sorted subpopulations, mock-treated vs infected vs bystander cells, were analyzed at early time-points of infection by RNA-sequencing (Hanna et al., 2019). When compared to mock-treated cells, the G1 to S transition related genes were up-regulated from 1 to 3 hpi in both bystander and infected cells, while they started to be down-regulated from 6 to 12 hpi, again in both cell subsets, with a more marked trend observed in infected cells (**Figure 7D**) likely underlining a delay in G1 to S transition. Again, similar trends were observed for the M to G1 transition related genes (**Suppl. Figure 5B**). When extended to later time-points (from 24 to 48hpi), the expression of these genes was more strongly up-regulated in bystanders than mock-treated cells, or infected cells in which the situation appears “frozen” (**Suppl. Figure 5C-D).** Overall, this suggests that mycobacterial shed material impacts the *D. discoideum* cell cycle, mainly delaying its progression through the G1 to S transition during the earliest phase of the infection, but this effect is relatively transient and does not represent a full block. In addition, the RNA-seq signatures are intermediate for bystander cells compared to mock-treated and infected cells (**Suppl. Figure 6A-B**), corroborating the measured interdivision times.

**Figure 7.**
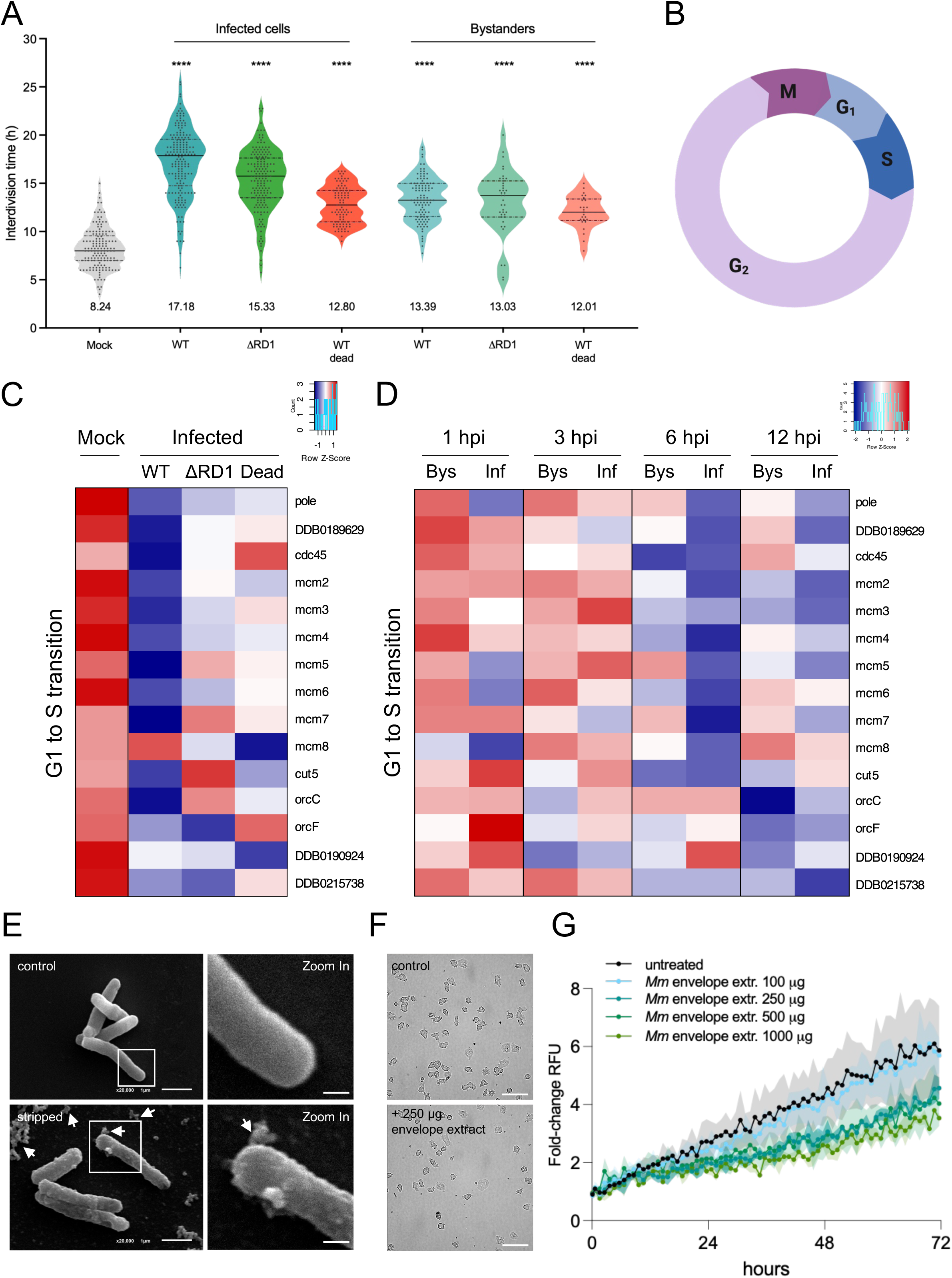
Cell cycle progression is delayed in bystander cells. (**A**) Single-cell analysis of the different infection cellular subsets (Mock, infected or bystander cells) by microfluidic shows a prolonged interdivision time in infected and bystander cells in *M. marinum* WT, ΔRD1 or WT dead infection contexts. Scattered violin plots shows total population values, with median (line) and interquartile (dashed line). p-values ****<0.0001, Tukey’s multiple comparison test. (**B**) *D. discoideum* cell cycle is divided in four phases: mitosis (M), cell growth (G1), DNA replication (S) and DNA-damage repair (G2). (**C**) Transcriptomic profiles of *D. discoideum* cells in contact with *M. marinum* WT, ΔRD1 or WT dead obtained by RNA-sequencing at 4hpi shows a down-regulation of the expression of most of the genes related to G1 to S cell cycle transition when compared to mock. (**D**) Transcriptomic profiles of *D. discoideum* infection cellular subsets (mock, infected and bystander cells) obtained by RNA-sequencing from samples collected at 1, 3, 6 and 12hpi shows an up-regulation in the expression of the G1 to S transition-related genes when compared to mock in bystander and infected cells from 1 to 3 hpi, switching to a down-regulation from 6 to 12 hpi. For panels **C** and **D**, read counts or logFC are represented, respectively, with row-normalized colored heatmap, with negative row z-score corresponding to blue colors and positive row z-score to red colors. Bys: Bystander cells, Inf: Infected cells. (**E**) Superficial stripping (white arrows) from *M. marinum* surface induces a rough surface but preserves bacilli structure, as seen by scanning electron microscopy (SEM). Scale bars = 1 µm, and 0.1 µm for zoomed in images. (**F**) *D. discoideum* conditioning with 250 µg of *M. marinum* envelope extract delays cell division after 24h, as observed by spinning-disc confocal microscopy (**G**). *D. discoideum* multiplication when conditioned with increasing quantities of *M. marinum* envelope extracts was assessed by plate reader over 72h, using a strain expressing a cytosolic mCherry reporter. Results are shown as fold-change of the fluorescence intensity (RFU) per well according to the first time-point, with mean +/-SD. Here is one representative of two independent biological replicates.

To attempt at recapitulating the impact of the envelope material on bystander cell cycle progression, we designed an “infection-free” assay. The most superficial layer of the mycobacteria envelope was gently stripped from *M. marinum* grown in broth, without profoundly altering the *M. marinum* morphology and surface structure (**Figure 7E**). After washing and concentrating, this fraction was used to condition *D. discoideum* cells. Incubation of 10^5^ cells with increasing concentration from 0.5, 1.25, 2.5 to 5 mg.ml^-1^ of envelope extract for 24 h, revealed a 30% decrease in growth compared to untreated cells (**Figure 7F-G**), validating the direct impact of cell wall components on host cell biology.

### The shed material induces a damage-response in bystanders, priming them to better resist subsequent infection

Starting minutes after their uptake and during their intracellular life, virulent mycobacteria damage the membrane of their compartment to establish their replicative niche and ultimately escape to the cytosol to further grow and disseminate to naïve cells. Host cells respond to MCV damage by recruiting the ESCRT and autophagy machineries to repair the membrane and contain bacteria (reviewed in Guallar-Garrido & Soldati, 2024). This damage-response is also perceptible at the transcriptomic level and is a hallmark of a mycobacteria intracellular infection. In this context, we compared the transcriptomic profiles of the ESCRT- and autophagy-related genes, between the two cell subsets of the infection during the first 12 hpi. A strong up-regulation was observed at 1 hpi, then progressively decreasing until 12 hpi in infected cells compared to mock-treated cells for both ESCRT- (**Figure 8A**) and autophagy- (**Figure 8B**) related genes. An intermediate profile was observed in bystander cells when compared to mock-treated cells, with a strong up-regulation being mostly observed at 1 hpi, underlying a transient induction of the damage response in these cells. Live microscopy images obtained during infection at 1.5 hpi reflected this induction. *D. discoideum* cells expressing GFP-Vps32 (as a reporter for the ESCRT machinery) or GFP-Atg8a (as a reporter for autophagy-mediated repair) were infected with mCherry-expressing *M. marinum*. As shown in **Figure 8C-D**, GFP-Vps32 and GFP-Atg8a positive structures are not only seen in the vicinity of bacteria in infected cells but accumulated as well in some bystander cells. While heterogenous, this phenomenon shows that a damage response can be induced in non-infected cells. To further investigate whether this is due to shed envelope material, GFP-Vps32 expressing cells were conditioned with an extract derived from *M. marinum* and monitored by high-content microscopy (**Figure 8E, Suppl. Figure 7A-B-C**). At 6 h post-conditioning, the median number of reacting cells per image (i.e. cells with at least one GFP-Vps32 positive dot) went up to 40% with 1.25 mg.ml^-1^ of envelope extract (against 22% for untreated cells) (**Figure 8F**). This increased reactivity was also reflected in a higher dot intensity (**Figure 8G**) and total area (**Figure 8H**). Interestingly, extracts derived from *M. marinum* WT or ΔRD1 induced a similar reaction, supporting an Esx-1-independent damage induction (**Suppl. Figure 7D**). The impact of this cellular conditioning on the fate of the infection was finally monitored by plate-reader. Conditioned cells were infected with *M. marinum* expressing a luminescent reporter and luminescence was quantitated over 72 hpi, as a proxy for intracellular growth over more than one infection cycle. A constant conditioning induced a drastic reduction of intracellular growth, resulting in a 3-fold reduction of the intracellular mycobacterial load (**Figure 8I**). When cells were transiently pre-conditioned for 6 h before infection, a milder intracellular growth defect was observed, with a 25% reduction. Altogether, these results demonstrate that cell conditioning with a mycobacteria envelope extract primes the cells to become more resistant to the infection.

**Figure 8.**
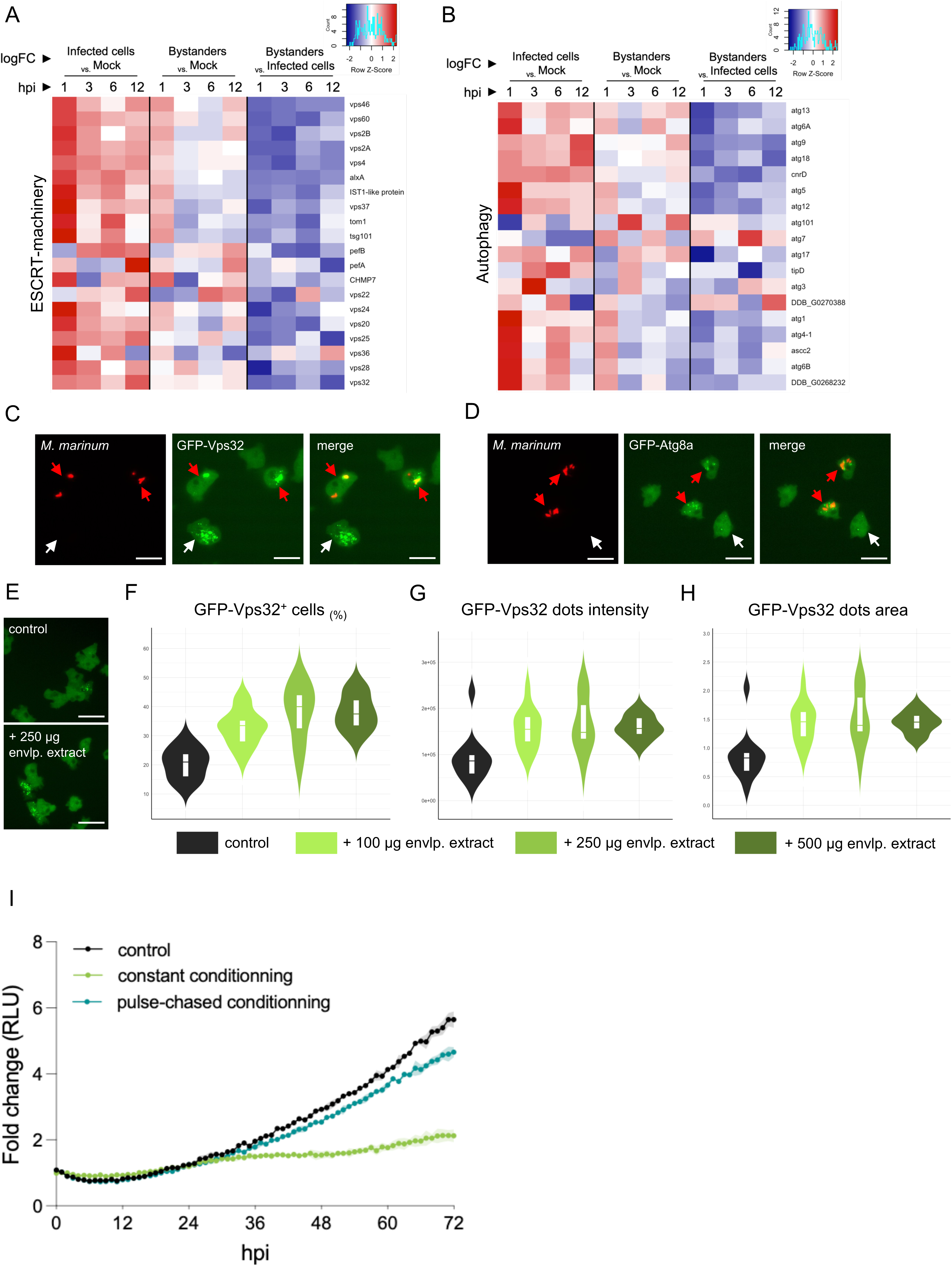
The membrane damage-response observed in bystanders during *M. marinum* infection can be recapitulated with *M. marinum* envelope extract. Transcriptomic profiles of *D. discoideum* infection cellular subsets comparisons (log Fold-change, logFC), obtained by RNA-sequencing from samples collected at 1, 3, 6 and 12hpi, shows transient up-regulation of (**A**) ESCRT-machinery or (**B**) autophagy related gene expression at 1hpi in bystander cells when compared to mock. While most of the genes are mostly down-regulated when compared to infected cells (reflecting lower up-regulation when compared to mock), few bystander-specific signatures can be observed, such as transient up-regulation in the expression of *atg7* at 1, 6 and 12hpi, or strong up-regulation of *DDB_G0270388* at 12hpi. LogFC of read counts are represented with row-normalized colored heatmap, with negative row z-score corresponding to blue colors and positive row z-score to red colors. (**C**) GFP-Vps32 positive bright structures accumulation is seen in infected (red arrows) and bystanders (white arrow) during *D. discoideum* infection with *M. marinum* pCherry. Scale bars = 10 µm. (**D**) GFP-Atg8a positive dots is seen in infected (red arrows) and bystanders (white arrow) during *D. discoideum* infection with *M. marinum* pCherry. Scale bars = 10 µm. (**E**) *D. discoideum* cell conditioning with *M. marinum* envelope extract for 6h induces similar GFP-Vps32 positive bright structures accumulation in some cells (Scale bars = 15 µm), as reflected by a dose-dependent significative induction of (**F**) the proportion of GFP-Vps32^+^ cells (cells with at least one GFP-Vps32 positive dot), (**G**) GFP-Vps32 dots fluorescence intensity (RFU) and (**H**) the GFP-Vps32 dots area. GFP-Vps32^+^ cells, dots intensity and area were measured by high-content microscopy, normalized per cell, and are represented as violin plots with boxplots for control or conditioned cells with 100, 250 and 500 µg of envelope extract. Results are representative of two independent experiments. (**I**) *D. discoideum* intracellular growth of *M. marinum* WT (expressing lux reporter) over 72 hpi is reduced when cells were pre-conditioned with *M. marinum* envelope extract, with a more drastic effect when constant conditioning is applied. Results (Fold-change in relative luminescence units, RLU) are representative of two independent experiments.

## Discussion

The surface of the mycobacterial cell wall is known to be labile and responsive to environmental changes and stresses. Although shedding of surface components has been reported in some intracellular *Mycobacterium* species, little was known about the fine dynamics of this process and its direct consequences on host cells. In this study, using *D. discoideum* as a simple cellular model to investigate the early stages of phagocytosis, we showed that the envelope of *M. marinum* is rapidly stripped from the bacterial surface upon internalization, accumulating in the phagosomal lumen. Interestingly, we and others (Beatty et al., 2002), detected the presence of multilamellar bodies (MLBs) in the phagosomal lumen early during infection (**Suppl. Figure 1H**). MLBs, consisting of multiple concentric layers of lipid-rich membranes, have been proposed to reflect the presence of undigested bacterial material (Cooper et al., 1986; Hohl, 1965). Thus, the presence of MLBs in the mycobacteria-containing vacuole likely reflects the accumulation of a large amount of shed lipid-rich mycobacterial cell wall components.

The early removal of surface components from mycobacteria was confirmed in a macrophage model and shown to be a conserved feature across a broad range of both pathogenic and non-pathogenic mycobacteria. Interestingly, prior work demonstrated that heat-killed *M. bovis* BCG can also release fluorescently labeled surface glycoconjugates (Rhoades et al., 2003). In our model, we showed that shedding rates were similar from live or dead bacteria (**Suppl. Figure 3A-B**). Taken together, these findings strongly suggest that the release of surface materials occurs through a passive, mechanical process, rather than an active, bacteria-driven mechanism. In line with this, high membrane trafficking observed shortly after mycobacteria uptake (Schnettger et al., 2017) might be sufficient to disrupt loosely associated surface components and distribute them throughout the connected endosomal compartment. Such mechanical stripping may represent a host-driven strategy to expose microbial pathogen-associated molecular patterns (PAMPs), as *M. marinum* bacilli that had lost their labeled surface components were more frequently ubiquitinated (**Figure 4D**).

Previous studies have documented the dissemination of mycobacterial components from infected to uninfected (bystander) cells, particularly focusing on surface-exposed glycolipids. First studies showed that shed labeled glycoconjugates traffic through the endosomal pathway, ultimately being released via extracellular vesicles (EVs), and later appearing in bystander cells (Beatty et al., 2002; Beatty & Russell, 2000). Similarly, Van den Elzen et al. described a mechanism by which bacterial glycolipids are secreted, in complex with the apolipoprotein E from infected macrophages and efficiently taken up by dendritic cells (van den Elzen et al., 2005). Such findings have shaped our understanding of mycobacterial lipid dissemination as a mechanism for antigen sharing and immune modulation. However, our observations suggest more than simple dissemination: a substantial accumulation of mycobacterial shed material occurs in bystander cells, as reflected by increasing levels of shed NHS-labeled proteins detected in these cells (**Figure 5**).

The mechanisms by which mycobacterial surface components are released from infected cells and reach bystander cells remained incompletely understood. Previous models have proposed the role of extracellular vesicles, either bacterial and/or host derived (Layre, 2020). Beatty and Russell (Beatty & Russell, 2000) speculated that mycobacterial products might be packaged into vesicles and released extracellularly, thus allowing them to “stimulate” bystander cells. *M. tuberculosis*-infected macrophages have been shown to release EVs containing both host and bacterial constituents, including LAM and LM (Athman et al., 2015). Interestingly, while the authors proposed that LAM release is dependent on mycobacteria viability, their own data show that substantial LAM release persists when cells are infected with γ-irradiated *M. tuberculosis*, supporting once again the passive release of mycobacteria surface components. While these studies provide important insights, most have focused on glycolipid components and at later stages post-infection. In contrast, our study provides the first quantitative demonstration that the release and accumulation in bystander cells of mycobacterial surface components, including proteins, occurs rapidly after phagocytosis initiation. This finding highlights how early, and efficiently, bacterial material can be exported and accumulate in surrounding cells. The specific mechanisms involved remain to be fully resolved, but our data suggest that a form of ESCRT-dependent exocytosis is required, as Δalix host cells failed to disseminate *M. marinum* shed material. In this model, while autophagy-dependent mechanisms seem to participate only partially to the trafficking and transfer of *M. marinum* shed material, the ESCRT machinery might be crucial to re-capture the cytosolic shed material in multivesicular bodies (MVB) and to regurgitate it for dissemination to neighboring cells.

While the dissemination of mycobacterial surface components to bystanders has been increasingly recognized, the functional consequences of this transfer, particularly at early time points, have remained largely unexplored. Here, we show that shed material accumulates in bystander cells within minutes of a primary infection of neighboring host cells and triggers a transient cellular response similar to the one observed in infected cells, characterized by an induction of membrane damage-response and a delay in the G1/S cell cycle transition. Both *M. tuberculosis* and *M. bovis* BCG have previously been shown to induce G_₀_/G_₁_ cell cycle arrest, apoptosis, and transcriptional reprogramming in infected cells (Cumming et al., 2017; Lai et al., 2007). Importantly, several purified mycobacterial lipids, such as MAME, PDIM, PIM_1,2_ and SL-1 are sufficient to trigger G_₁_/S arrest, even in the absence of live bacteria (Cumming et al., 2017). This aligns with our observation that exposure to *M. marinum* envelope material alone, in the absence of active infection, can induce a delay in cell cycle progression. Interestingly, *in vivo* studies have shown that while *M. tuberculosis* infected cells undergo cell cycle arrest, bystanders are locally proliferating in lung, contributing to tissue remodeling and immune recruitment (Huang et al., 2018). Consistent with this, we observed an up-regulation of genes involved in G_1_/S transition in bystanders at later stages (**Suppl. Figure 4**). This suggests a dual response: bystander cells may transiently pause their cell cycle upon sensing shed material but remain functionally competent and potentially better equipped to control infection upon subsequent bacilli uptake.

Several mycobacterial surface components are known to possess immunomodulatory properties that could explain such priming effects. Notably, virulence-associated lipids such as PDIM, PGL, and SL-1 have been individually proposed to be implicated in immune evasion. For example, expression of *M. leprae* PGL-1 in *M. bovis* BCG promotes uptake by dendritic cells while dampening inflammatory gene expression (Tabouret et al., 2010), while *M. tuberculosis* SL-1 has been shown to enhance lysosomal biogenesis and exocytosis (Umar et al., 2025). The use of other purified lipids, including MAME, FAME, and trehalose dimycolate (TDM), can induce phenotypes associated with lysosomal storage disorders or alter host membrane properties and fluidity (Fineran et al., 2016; Mishra et al., 2020; Srivatsav & Kapoor, 2023). In our study, we showed that PGL was found among the shed and disseminating material. The fact that the damage response was observed in bystander cells, independently of the presence of the major damaging factor EsxA, indicate that other mycobacterial surface components, such as PGL, may contribute at high concentrations to the induction of a membrane damage response and help bystanders sense and respond to the infection. Following this line of thoughts, bystanders, though not initially infected, can become targets during secondary or tertiary infection cycles following the egress of bacilli from primary infected cells. Early exposure to shed bacterial components may thus serve to sensitize these cells, enhancing their ability to recognize and resist the infection at later stages. Taken together, these observations suggest that mycobacterial surface components, once shed, act not only as virulence factors within infected cells but also as modulators of bystander cell biology. Early exposure appears to reprogram bystanders toward a more alert, resistant state, a form of innate immune "preconditioning" that may shape the outcome of infection spread at the population level.

## Acknowledgments

We gratefully acknowledge Dr. Dimitri Moreau, Dr. Stefania Vossio and Dr. Vincent Mercier, from the ACCESS Geneva Imaging Facility (University of Geneva) for their assistance and advice with high-content microscopy imaging and segmentations. A special thanks to Jérôme Bosset, from the Photonic Bioimaging Center (University of Geneva) for his help in high-resolution microscopy and SEM images acquisition. We acknowledge Dr. Claire Mulholland and Dr. Michael Berney, from Albert Einstein College of Medicine, for kindly providing us *M. tuberculosis* BSL-2 strains and advising us on their cultivation protocols. We thank Céline Michard for kindly providing us the *M. marinum* Δpks15/1 mutant, as well as the *D. discoideum* Δalix cell line. We thank Manon Mottet for having performed the microfluidic experiments. We acknowledge the receipt of reagents and bacteria species from BEI Resources, NIAID, NIH, including purified PGL and PDIM, as well as all non- *M. marinum* mycobacteria species used in this work. We thank Dr. Aleksandar Salim (from Dr. Sascha Hoogendoorn lab, University of Geneva) for the synthesis of the azido *p*HB precursor for PGL metabolic labeling. This work was supported by Swiss National Science Foundation grants 310030_188813 and 310030_219364, and an EMBO post-doctoral fellowship earned by MF (ALTF 715-2021).

## Author contributions

All authors made substantial contribution for this study, and approved accuracy and integrity of the work presented in this manuscript. DD, MF and TS contributed equally to all conceptual aspects of the project. TS acquired fundings required for this project. DD and MF both worked on all experimental designs, acquisitions, analyses, and interpretations. LR participated in all transcriptomic analyses. MF and TS wrote and revised the manuscript.

## Material and methods

### *D. discoideum* and BV-2 cell culture

The *D. discoideum* axenic strain Ax2(ka) was maintained in culture at 22°C in 10 cm culture dishes with HL5c medium including glucose supplemented with vitamins and microelements (ForMedium, ref HLE2), supplemented with 100 U/mL of penicillin and 100 µg/mL of streptomycin (Gibco, ref 15140-122). Plasmids were transfected into *D. discoideum* by electroporation for integration of fluorescent reporters at the safe haven act5 locus (Paschke et al., 2019) and selected with 50 µg/mL of hygromycin. Plasmids and resulting cell lines are listed in **Supplementary table S1**. BV-2 microglial cells were grown at 37°C, 5% CO_2_ in DMEM high glucose (Sigma-Aldrich, ref D5796-500ML) complemented with 10% Fetal Bovine Serum (Gibco, ref 10270-106), supplemented with 100 U/mL of penicillin and 100 µg/mL of streptomycin as well.

### Mycobacteria strains, plasmids and culture

All mycobacterial strains and species used in this study are listed in **Supplementary table S1**. *M. marinum* M strain was grown in Middlebrook 7H9 (BD Difco, ref 271310) enriched with 10% OADC (Oleic acid Albumin Dextrose Catalase, Becton Dickinson, ref 212351), 0.2% glycerol (Sigma-aldrich, ref G2025-1L) and 0.05% tyloxapol (Sigma-aldrich, ref T8761-50G) at 32°C, in flasks with shaking at 150 rotation per minute (rpm) and in the presence of 5 mm glass beads to minimize bacteria clumping. Other mycobacteria strains were cultured similarly, but at 37°C, except for *M. abscessus* which was grown at 32°C *M. marinum* WT or ΔRD1 expressing cytosolic GFP (pmsp12::GFP for WT, pGFPHYG2 for ΔRD1), pCherry (pMV306::pCherry10) or a bacterial luciferase (pMV306::lux) were used and selected with kanamycin 50 µg/mL or hygromycin 100 µg/mL as appropriate. The BSL-2 approved *M. tuberculosis* mc^2^6206 strain (genotype H37Rv ΔpanCD ΔleuCD), kindly provided by Prof. Berney, was cultured in 7H9 enriched with 10% OADC, 0.2% glycerol, 0.05% tyloxapol and supplemented with 24 µg/mL pantothenate (Thermoscientific Acros, ref 24330-0050) and 50 µg/mL L-leucine (Sigma-aldrich, ref L-8912) (Mulholland et al., 2024).

### Mycobacteria cell wall labeling

- **Labeling of surface-exposed carbohydrates** Mycobacteria were cultured to reach an optical density (OD_600_) ∼0.8-1 (i.e. mid-log phase, reaching a biomass of around 1.3 to 1.8 x 10^8^ bacteria/mL). 1 x 10^9^ bacteria were centrifuged at 3,000 rpm for 10 min, at room temperature (RT). Resulting pellet was washed 3-times with Phosphate Buffer Saline (PBS), and re-suspended with 500 µL of 0.1M sodium acetate, pH 5.5 containing 1mM sodium periodate to oxidize hydroxyl terminal groups and form reactive aldehyde residues. After an incubation of 20 min at 4°C with gentle rotation, 0.1mM glycerol was added to stop the reaction. Bacteria suspension was then washed 3-times with PBS, re-suspended in 500 µL PBS containing 1 mM fluorescent-labeled hydrazide (Alexa Fluor^TM^ 488-hydrazide, Invitrogen ref A10436) and incubated for 1h at RT, in the dark with shaking. After 3 PBS-washes, pellet was finally re-suspended in PBS, 7H9 or HL5c for infection.
- **Labeling of surface-exposed proteins** Mycobacteria were cultured as for surface-exposed carbohydrates labeling. 1 x 10^8^ bacteria were centrifuged at 3,000 rpm for 10 min, at RT. Pellet was washed 3-times with PBS-containing 0.2M sodium bicarbonate pH 8.8, re-suspended in 500 µL PBS-0.2M sodium bicarbonate containing 1 mM fluorescent succinimidyl/NHS ester (Alexa Fluor^TM^ 488-NHS, Invitrogen ref A20000 - Alexa Fluor^TM^ 594-NHS, Invitrogen ref A20004) and incubated for 1h at RT, in the dark with shaking. After 3 PBS-washes, pellet was finally re-suspended in PBS, 7H9 or HL5c for infection.
- **Metabolic labeling of peptidoglycan** Metabolic labeling of peptidoglycan was coupled with the two previous labeling protocols. Briefly, before incubation with fluorescent dyes (hydrazide or NHS), bacterial suspensions were washed 3-times with 7H9 and incubated for 1h with 500µM HADA (7-Hydroxycoumarin-3-carboxylic-acid-D-Alanine, Tocris Bioscience ref 6647/5) or 25µM NADA (3-[7-nitrobenzofurazan]-carboxamide-D-Alanine, Tocris Bioscience ref 6648/5) at 32°C in the dark with shaking. After 3 washes in PBS, suspensions were processed for surface-exposed carbohydrates or proteins labeling.
- **Metabolic labeling of PGL** Protocol for metabolic labeling of *M. marinum* PGL was adapted from Guzmán et al., 2024. *M. marinum* were cultured in 7H9 containing 500µM 3-azido p-hydroxy benzoic acid (*p*HB) for 18h at 32°C with shaking, to reach an OD_600_ ∼0.8-1. After incubation, 1 x 10^8^ bacteria were pelleted at 3,000 rpm, 10 min RT. Resulting pellet was washed 3-times with PBS and labeled with 30µM DBCO-AF488 (BaseClick ref BCFA-237) for 1h at RT, in the dark with shaking. Bacteria were then washed 3-times with PBS and resulting pellet was re-suspended in PBS, 7H9 or HL5c for infection.

### Flow cytometry assay

Cell wall labeling efficiency was quantitatively measured by flow cytometry. After labeling protocol, bacteria re-suspended in PBS were adjusted to a final OD_600_ ∼0.1. Samples were analyzed using a SONY SH800 flow cytometer and data were processed with the free web-based application Floreada.io.

### Colony forming units (CFU) counting

Impact of cell wall labeling protocols on bacterial viability was evaluated by CFU counting. After labeling protocol, bacteria re-suspended in 7H9 were serially diluted to 10^-4^ and 100µL of each dilution was plated onto Middlebrook 7H11 plates (Difco, ref 212203). Plates were incubated at 32°C for 1 week, until colony forming.

### Total lipid extraction and thin-Layer chromatography (TLC)

PGL effective labeling was controlled by analysis of total lipid extract from labeled *M. marinum* suspensions. After labeling protocol, bacteria pellets were snap-frozen, and Bligh and Dyer extraction was done as previously described (Bligh & Dyer, 1959). After lipid extraction, lipid samples were re-suspended in 100µL chloroform and spotted on silica High performance TLC plates (Merck Millipore, ref Z740222). Glycolipids were separated in a system containing chloroform:methanol (60:40, v:v), and PDIM and neutral lipids were separated with a system consisting of petroleum ether:diethylether (90:10, v:v). Both systems ran to 1cm below the upper border. After drying plates under a fume hood, fluorescent lipids were visualized with UV illumination (Fusion Fx device, Vilber Lourmat) and total lipids were detected by charring with a solution containing 0.63g MnCl_2_.4H_2_O, 60mL water, 60mL methanol and 4mL concentrated sulfuric acid). Lipid standards were kindly obtained from BEI Resources: purified PGL from *M. canettii* (NR-36510) and purified PDIM from *M. tuberculosis* (NR-20328).

### *M. marinum* envelope preparation

Protocol for envelope extraction from *M. marinum* surface was adapted from a protocol previously developed to extract extracellular vesicles embedded in the extracellular matrix produced by *M. ulcerans* (Foulon et al., 2022). Both fractions are labile, i.e. weakly attached to the myco-membrane, and can be removed by gentle mechanical disruption. Accordingly, *M. marinum* was cultured in 80 mL 7H9 without beads, to reach an OD_600_ ∼0.8-1. Bacteria were centrifuged at 3,000 rpm for 10 min, at RT and pellet was washed 2-times with PBS to remove secreted particles not attached to the envelope/cell wall. After washing, 10 glass beads (5 mm diameter) were added to the pellet with 1 mL cold water, or HL5c, and 4 cycles of 45 sec vortexing were applied (with 1 min resting on ice in between). The suspension was then centrifuged 5 min at 3,000 rpm, RT, and supernatant was collected and filtered on 0.45µm filter units to obtain the capsular extract (approximatively 10 to 15 mg for 80 mL culture). Envelope extracts were lyophilized for long-term storage or directly used to condition *D. discoideum* cells.

### Scanning electron microscopy (SEM)

To image the bacterial surface before and after envelope extraction, 1 x 10^8^ bacteria were seeded on poly-L-lysine (1 mg/mL) coated glass coverslips in a 24-well plate. After a brief centrifugation at 600 rpm, coverslips were incubated for 30 min to improve bacteria adhesion, then fixed with 4% paraformaldehyde (Fluka, ref 76240) plus 2% glutaraldehyde (Electron Microscopy Sciences, ref 16120) in PBS overnight at RT in the dark. Samples were then post-fixed with 1% osmium tetroxide (Electron Microscopy Sciences, ref 19190) in 0.1M sodium cacodylate for 1h in the dark. After rinsing 3-times with 0.1M sodium cacodylate (Electron Microscopy Sciences, ref 12310), coverslips were dehydrated in a graded ethanol series (from 30 to 100% ethanol, 5 min incubation each and repeating the incubation in ethanol 100% twice). Ethanol was them replaced with hexamethyldisilazane (HMDS, Milipore ref 8043241000) and coverslips were left to dry overnight, before to be mounted on carbon coated stubs, coated with 5 nm gold-palladium, and imaged on a JEOL SEM (JSM-6510LV) at 30 kV.

### Infection assay

In *D. discoideum* cells, infections were performed as previously described (Hagedorn & Soldati. 2007), with few modifications. Mycobacteria suspensions were washed but not disrupted by syringing, to avoid envelope removal. After infection at a multiplicity of infection (MOI) of 10 for *M. marinum* WT and MOI 20 for *M. marinum* ΔRD1, cells were left for 20 min of phagocytosis, extracellular bacteria were washed off with 3 to 5 washes with 5 mL HL5c, and infected cells were re-suspended in filtered HL5c containing a bacteriostatic dose of 5 U/mL of penicillin and 5 µg/mL streptomycin, to prevent extracellular growth of *M. marinum*. Mock cells were treated similarly without adding bacteria. Time post infection was defined relative to the moment when the bacteria were added to the attached cells. For BV-2 cells, the protocol of infection was followed similarly, with slight adaptations: 5 x 10^6^ cells were plated in DMEM complemented with FBS without antibiotics the day before the infection, and were infected with an MOI of 5 at 32°C.

### Plate reader assay

*D. discoideum* cells infected with *M. marinum* pMV306::lux were seeded (2 x 10^5^ cells per well) in a white microwell^TM^ 96-well plate (Nunc, ref 136101) and sealed with a gas permeable moisture barrier seal (4titude, ref 4ti-0516/96). Bioluminescence intensity, measured in relative light units (RLU) as a readout for bacterial growth, was measured every 1h for 72h, at a constant temperature of 25°C using a Synergy Mx microplate reader (Biotek).

### Live confocal microscopy

After infection, cells were plated in a µ-dish IBIDI 35 mm (ref 81156) for point-scan confocal high-resolution microscopy or in a 96-well IBIDI dish (ref 89626) for high-content spinning disc microscopy. For high-resolution live microscopy, images were recorded with a Leica Stellaris 8 63×1.4 NA oil immersion objective. For high-content live microscopy, images were recorded with a 60x water immersion objective mounted on the ImageXpress Micro XL HC microscope. All segmentations and quantifications were done with the MetaXpress^R^ molecular device software.

### Immunofluorescence staining

For immunofluorescence staining, infected *D. discoideum* cells were fixed 2hpi with ultra-cold methanol as previously described (Hagedorn et al., 2006), to achieve optimal preservation and permeabilization of the cells. After fixation, samples were blocked with PBS containing 0.2% gelatin (PBG) for 1h at RT. After 3 rinsing, cells were incubated for 1h with primary antibodies directed against p80 (Lima & Cosson. 2019) or ubiquitin FK2 (Enzo Life Sciences, ref BML-PW8810-0500) diluted at 1:10 or 1:1000, respectively, in PBG. Cells were rinsing once again 3-times and incubated with secondary antibodies (goat anti-rabbit or goat anti-mouse, both coupled with Cy5) diluted at 1:1000 in PBG, mixed with DAPI, diluted 1:100, to mark nuclei. After final rinsing, coverslips were carefully mounted on Prolong Gold Antifade Mountant (Invitrogen, ref P36934) and left to dry overnight before imaging. Images were recorded with a Leica Stellaris 8 63×1.4 NA oil immersion objective.

### Cell conditioning

*D. discoideum* cells (WT or expressing GFP-Vps32) were conditioned in 96-well black plates with transparent bottom (IBIDI, ref 89626). 1.25 x 10^4^ (for cell multiplication assay) or 5 x 10^4^ (for cell damaging assay) cells were seeded per well in filtered HL5c and incubated at 22°C half an hour for cell adhering. Medium was then replaced by envelope extract solubilized in filtered HL5c (conditioning medium) and images were recorded every 30 min or 1h for 6-12h using a 60x water immersion objective mounted on the ImageXpress Micro XL HC microscope.

### Dissemination assay

This assay was adapted from the dissemination assay previously developed to quantitatively monitor cell-to-cell spreading of mycobacteria (Hagedorn et *al.* 2009). Here, *D. discoideum* expressing cytosolic act5::mCherry were infected as described above with *M. marinum* pCherry prelabeled with 488-NHS, constituting the “donor” population. This population was then added to *D. discoideum* not expressing any reporters, being the “receiver” population (already seeded in 96-well black plate with transparent bottom (IBIDI, ref 89626) at a donor:receiver ratio of 1:1. Images were recorded with a 60x water immersion objective mounted on the ImageXpress Micro XL HC microscope.

### Long-term single cell analyses by microfluidic

Protocol for long-term single cell imaging of *D. discoideum* was previously described in Mottet et *al.* 2021. Briefly, after infection, 3.5 x 10^6^ cells were added on micropatterned coverslips in a 35 mm dish, and the whole microfluidic device was carefully mounted and connected to a syringe containing filtered HL5c medium, loaded to a syringe pump maintaining a constant flow rate of 10µL/min. Images were recorded with a 63x glycerol objective mounted on a spinning disc confocal microscope (Intelligent Imaging Innovations Marianas SDC mounted on an inverted microscope (Leica DMIRE2)) at 25°C.

### RNA-sequencing

For RNA-sequencing performed on infected populations at 4hpi, *D. discoideum* cells infected with *M. marinum* WT, ΔRD1 or dead WT (killed with 10 µM rifabutin overnight) for 4h before to be washed and re-suspended in TRI-reagent (Sigma-aldrich, ref T9424) for RNA isolation. For cellular subset characterizations*, D. discoideum* cells infected with *M. marinum* GFP were sorted at specific time-points (1, 3, 6, 12, 24, 36 and 48 hpi) using an Astrios device (Beckman) at 4°C, based on GFP-positive (infected cells) or GFP-negative (bystander cells or mock) gating strategies. For each subset, 5 x 10^5^ cells were collected, centrifuged and re-suspended in TRI-reagent. RNA isolation was extracted using the Directzol RNA extraction kit (Zymo research, ref R2053) following manufacturer’s instructions. Quality of RNA librairies, sequencing and bioinformatic analysis were performed as previously described (Hanna et al., 2019).

### Statistical analyses

Data are presented as means and SD or SEM (standards deviations or standard errors of the mean) and were analyzed with GraphPad Prism 7.0 software (GraphPad Software, San Diego, CA, USA).

---

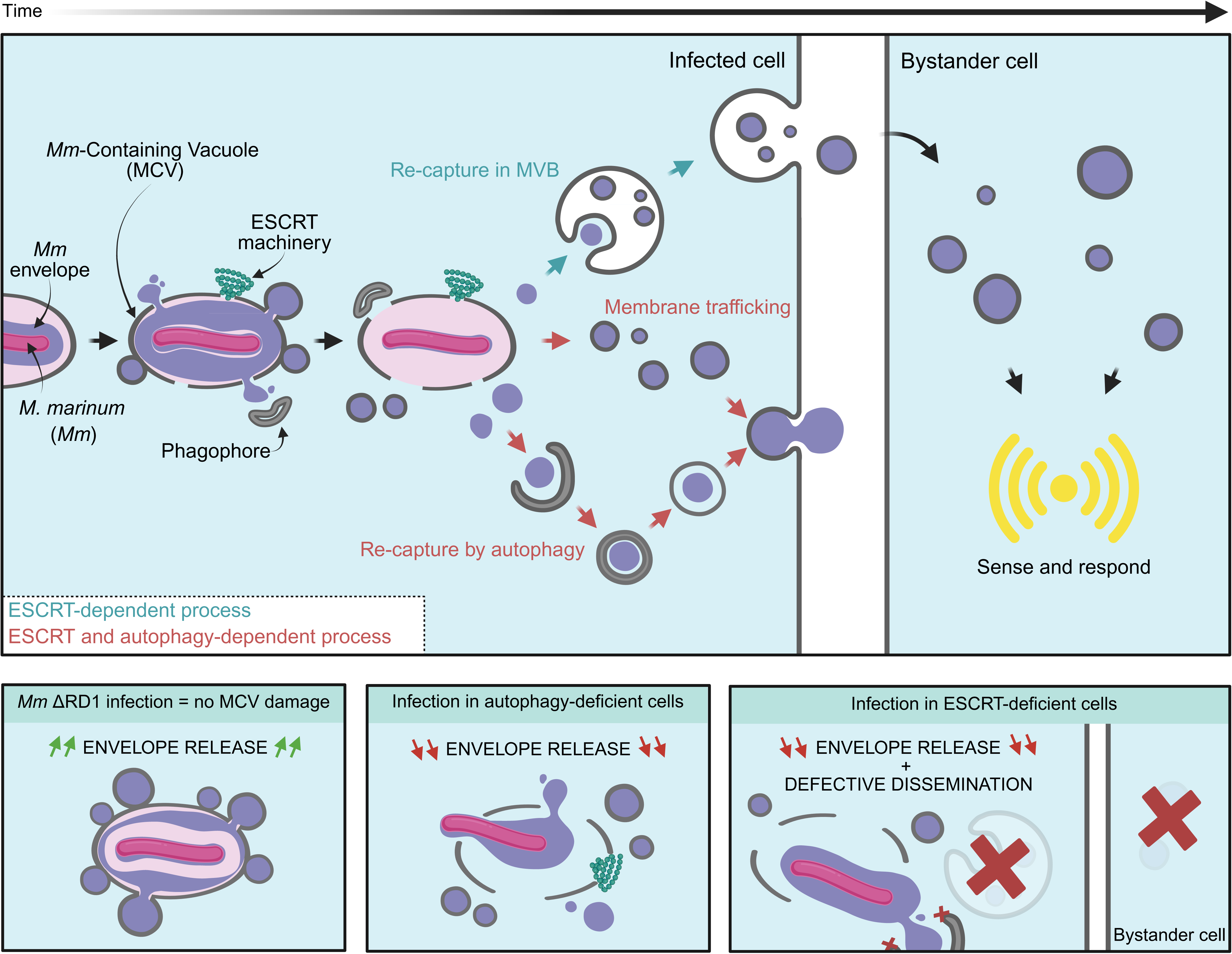

## Notes

### Competing Interest Statement

The authors have declared no competing interest.

### Summary of Updates

Figures legends missing in the main manuscript file have been added, and figures legends in the supplementary material file have been moved before the figures.

