## Supplementary material for "Mycobacterial surface shedding drives bystander cells response during early intracellular infection"

**Title**

**Affiliations**

**Supplementary Figures Legends**

**Supplementary Figure 1. *M. marinum* surface labeling is efficient and not affecting bacilli viability.** Fluorescence-Activated Cell Sorting (FACS) by flow cytometry shows a shift in fluorescence intensity median (expressed as RFU) for (**A**) 488-hydrazide or (**B**) 594-NHS labeled bacteria, when compared to unlabeled bacteria (control). Results are representative of three independent replicates. (**C**) CFU counting before and after labeling protocols shows no difference in *M. marinum* viability when compared to respective controls (subjected to the same steps than labeled bacteria, without the dyes). (**D**) Metabolic labeling of PGL, monitored thanks to copper-free clicked chemistry with 488-DBCO, shows effective incorporation of the azido-precursor *p*HB in the cell wall of *M. marinum* WT while it stays stuck inside the cytoplasm of *M. marinum* Δpsk15/1 (defective for PGL production) Scale bars = 0.5 µm. (**E**) TLC analysis on total lipid extract from pre-labeled *M. marinum* WT or Δpsk15/1 (defective to produce PGL) shows a band for metabolically labeled PGL (black arrowhead) only in *M. marinum* WT sample. No incorporation is visible in the close related lipid species PDIM (white arrowhead). PGL (and other glycolipids) were separated on a system consisting of a mixture of chloroform:methanol ($CHCl_3$:MetOH, 60:40, v:v) and PDIM (and neutral lipids) were separated with petroleum ether:diethylether (90:10, v:v). TLC plates were revealed for fluorescence, then charred with a sulfuric acid containing solution to see total lipids. (**F**) FACS analysis of *M. marinum* WT or Δpsk15/1 metabolically labeled for PGL shows a shift in fluorescence intensity only when *M. marinum* WT was provided with both the azido *p*HB precursor and 488-DBCO. No shift is observed in bacteria incubated only with 488-DBCO, or when the *M. marinum* Δpsk15/1 mutant is used. Results are representative of three independent replicates. (**G**) *M. marinum* (expressing cytoplasmic GFP reporter) is not replicating in the first 12hpi inside *D. discoideum*, when intracellular growth (reflected as fold-change in RFU signal) is followed by plate reader after cell infection. N=3 independent experiments, with n=3 technical replicates within each of them. (**H**) The presence of multilamellar bodies in the mycobacterium-containing compartment at 1.5hpi in *D. discoideum* cells, as seen with transmission electron microscopy (TEM) images. Scale bars = 0.6 µm and 0.2 µm for insets.

**Supplementary Figure 2.** ***M. marinum*** **ΔRD1 is labeled similarly to the WT strain for surface-exposed carbohydrates and proteins.** Labeling with 488-hydrazide (**A**) or 594-NHS (**B**) shows a homogenous surface labeling for both *M. marinum* WT and ΔRD1 by confocal microscopy. Scale bars = 2 µm and 0.5 µm in inset images for panel **A**, and 1 µm and 0.5 µm in inset images for panel **B**. FACS analysis shows a similar shift in fluorescence intensity (RFU) for *M. marinum* WT and ΔRD1 labeled with 488-hydrazide (**C**) and 594-NHS (**D**). Results are representative of three independent replicates. (**E**) CFU counting before and after labeling protocols shows no difference in *M. marinum* ΔRD1 viability when compared to respective controls (subjected to the same steps than labeled bacteria, without the dyes), and when compared to *M. marinum* WT (**Suppl. Figure S1**). (F) High-resolution live acquisition of *D. discoideum* infected cells, 10 min post-infection, shows similar early shedding and compartment filling for *M. marinum* WT and ΔRD1 (expressing cytoplasmic pCherry and labeled for surface components (488-NHS). Scale bars = 2 µm.

**Supplementary Figure 3.** ***M. marinum*** **envelope is shed even with dead bacteria, and shed material is trafficking among endocytic pathway in BV-2 infected cells. (A)** Quantification by high-content microscopy of 594-NHS labeling intensity shows a similar shedding of surface-exposed proteins when *D. discoideum* cells were infected with *M. marinum* WT alive (black) or killed with antibiotics (dark red). **(B**) Shedding rates were estimated based on the slopes of the curves from 0 to 4 and 4 to 12 hpi for each condition. (**C to D**) BV-2 cells infected with *M. marinum* WT pre-labeled with (**C**) 594-NHS or (**D**) 488-hydrazide are filled with small compartments filled with labeled components, dynamically spreading inside the cells throughout the time-course of the infection (from 2 to 12hpi). Images are shown as max projection of 5 z-stacks. Scale bars = 10 µm.

**Supplementary Figure 4.** ***M. marinum*** **488-hydrazide labeled shed material is actively trafficking and disseminating to bystander cells. (A)** In *D. discoideum* infection, large field of view shows accumulation of 488-hydrazide labeled shed material in non-infected cells (white arrows) at 2hpi. Scale bars = 20 µm. (**B**) High-content microscopy-based segmentation allows to measure shed material accumulation in the cell cytosol (yellow surface) by excluding the extended bacterial mask (blue surfaces). Total fluorescence intensity labeled *Mm* shed material (488-hydrazide) was measured in infected (red lines) and bystanders (grey lines) cells and normalized to the median in mock for *D. discoideum* WT (**C**), Δatg1 (**D**) or Δalix (**E**) cells infected with *M. marinum* WT. Results (median +/- SEM) are shown as representative of one to three independent experiments. (**F**) Trafficking rates in infected and bystander cells were calculated based on the first 3 hpi. (**G**) Dissemination ratios were calculated for each condition. Results are shown as bar plots +SD, with individual dots representing two independent biological replicates.

**Supplementary Figure 5. Transcriptomic profiles of *D. discoideum* infected populations shows several alterations in cell cycle progression.** (**A**) *D. discoideum* cells in contact with *M. marinum* WT, ΔRD1 or WT dead obtained by RNA-sequencing at 4hpi shows a down-regulation of the expression of most of the genes related to M to G1 cell cycle transition when compared to mock. (**B**) Transient signature in M to G1 transition is also observed in bystander and infected cells, with an up-regulation in gene expression from 1 to 3 hpi, followed by a do-regulation from 6 to 12 hpi when compared to mock. Read counts and logFC are represented with row-normalized colored heatmap, with negative row z-score corresponding to blue colors and positive row z-score to red colors. Transcriptomic profiles extended to later stages of the infection (24, 36 and 48hpi) shows an inverted trend, with higher up-regulation in the expression of most of the genes related to (**C**) G1 to S or (**D**) M to G1 transition in bystanders from 24 to 48hpi, while those genes stay down-regulated in infected cells. LogFC of read counts are represented with row-normalized colored heatmap, with negative row z-score corresponding to blue colors and positive row z-score to red colors. Bys: Bystander cells, Inf: Infected cells.

**Supplementary Figure 6. RNA-seq analyses reveal an intermediate transcriptomic signature in bystander cells.** Multidimensional Scaling (MDS) plots on pairwise dissimilarities shows clustering of samples by time-points (**A**) or by cellular subset (**B**). Each dot represents an individual sample, colored by time-point in (A) or by cellular subset in (B). Volcano plots of DEGs in (C) infected or (D) bystander cells compared to mock-treated cells shows an overall stronger response in infected cells at each time-point. All genes with an absolute log2 fold-change >0.058 and a false-discovery rate <0.05 were considered differentially expressed. Blue dots represent down- regulated genes and red dots represent up-regulated ones. Non-differentially expressed genes are colored in grey.

**Supplementary Figure 7. *D. discoideum* conditioning with *M. marinum* envelope extract induces damage-response (reflected by GFP-Vps32 recruitment) in an esx-1 independent manner.** (**A**) Percentages of reacting cells (GFP-Vps32+ cells), (B) GFP-Vps32 dots intensity and (**C**) area, normalized per cell and per image, all show an increase in cell conditioned with 100, 250 and 500 µg of *M. marinum* WT envelope extract, significatively different from control cells at 6h post-conditioning. (**D**) Both *M. marinum* WT and ΔRD1 envelope extracts induce similar reaction, as reflected with higher percentages of reacting cells (GFP-Vps32+ cells) at 6h conditioning when compared to control (untreated cells). Results are representative of at least two independent experiments.

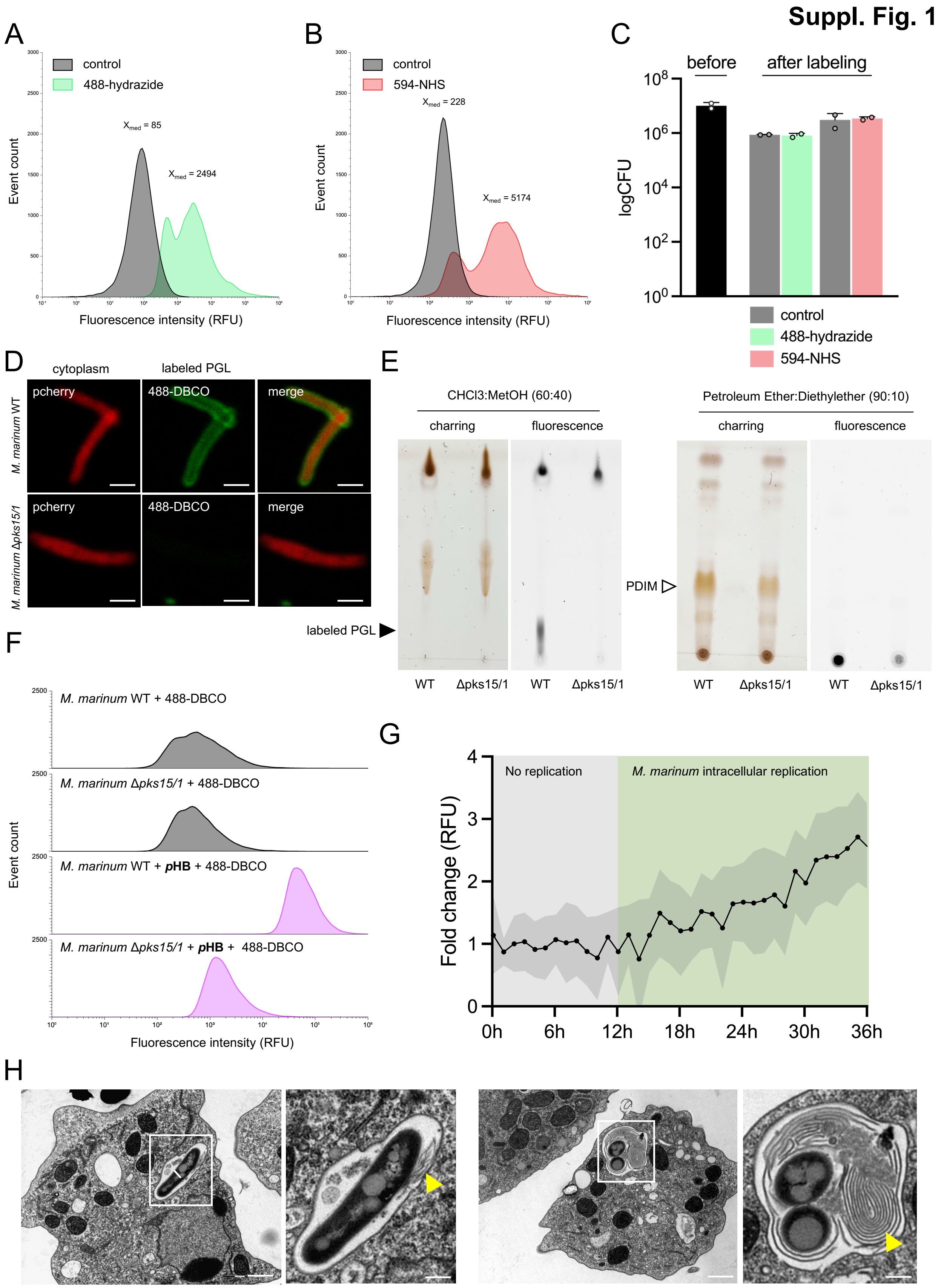
Suppl. Fig. 1
A
control
488-hydrazide
$X_{med}$ = 85
$X_{med}$ = 2494
Event count
Fluorescence intensity (RFU)
B
control
594-NHS
$X_{med}$ = 228
$X_{med}$ = 5174
Event count
Fluorescence intensity (RFU)
C
before
after labeling
logCFU
control
488-hydrazide
594-NHS
D
cytoplasm
labeled PGL
M. marinum WT
M. marinum Δpks15/1
pcherry
488-DBCO
merge
E
CHCl3:MetOH (60:40)
Petroleum Ether:Diethylether (90:10)
charring
fluorescence
labeled PGL
PDIM
WT
Δpks15/1
F
M. marinum WT + 488-DBCO
M. marinum Δpks15/1 + 488-DBCO
M. marinum WT + pHB + 488-DBCO
M. marinum Δpks15/1 + pHB + 488-DBCO
Event count
Fluorescence intensity (RFU)
G
No replication
M. marinum intracellular replication
Fold change (RFU)
0h
6h
12h
18h
24h
30h
36h
H

**Suppl. Fig. 2**

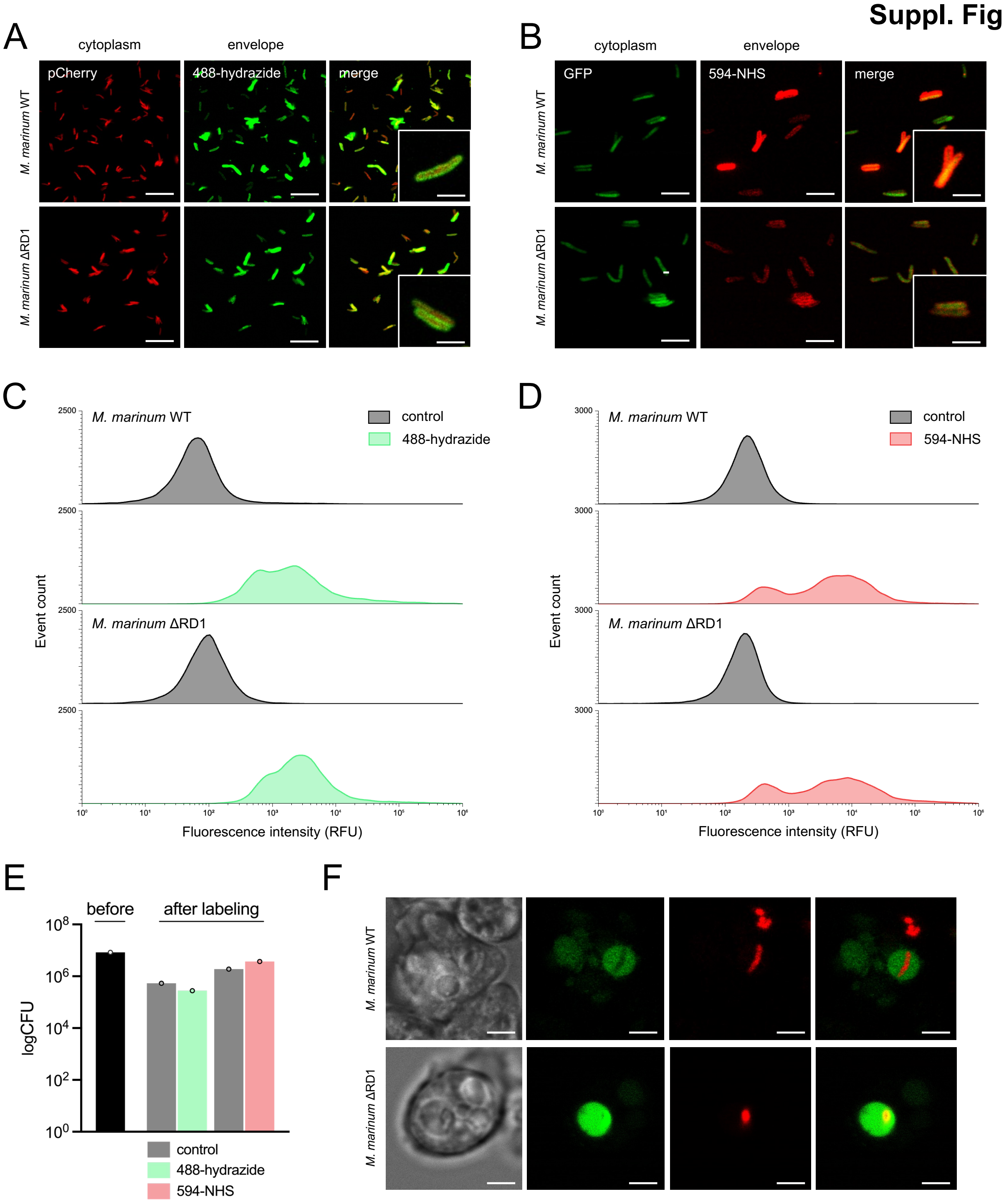
A
cytoplasm
envelope
pCherry
488-hydrazide
merge
M. marinum WT
M. marinum ΔRD1
B
cytoplasm
envelope
GFP
594-NHS
merge
M. marinum WT
M. marinum ΔRD1
C
M. marinum WT
M. marinum ΔRD1
control
488-hydrazide
Event count
Fluorescence intensity (RFU)
D
M. marinum WT
M. marinum ΔRD1
control
594-NHS
Event count
Fluorescence intensity (RFU)
E
before
after labeling
logCFU
control
488-hydrazide
594-NHS
F
M. marinum WT
M. marinum ΔRD1

### Suppl. Fig. 3

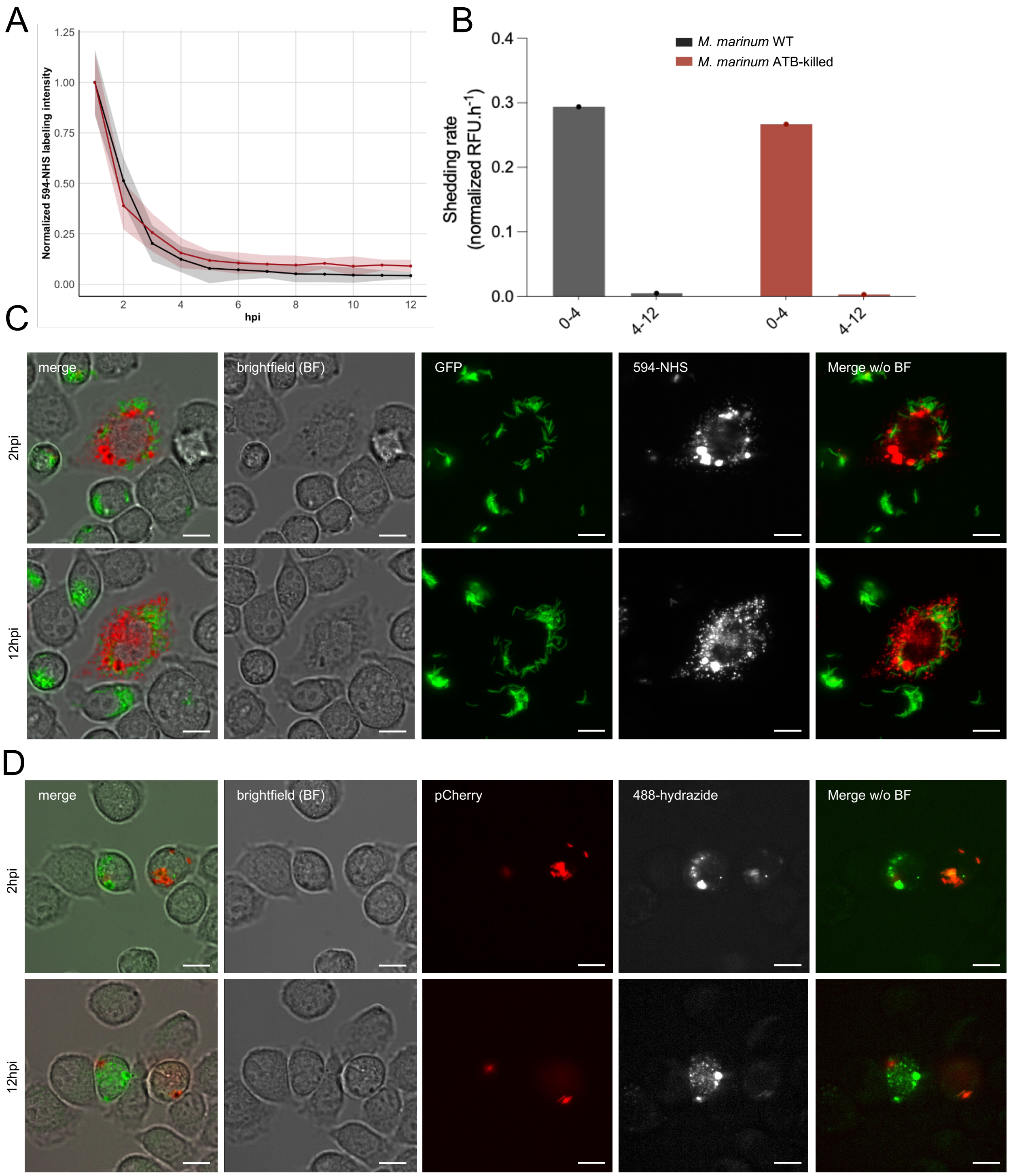


111

**Suppl. Fig. 4**

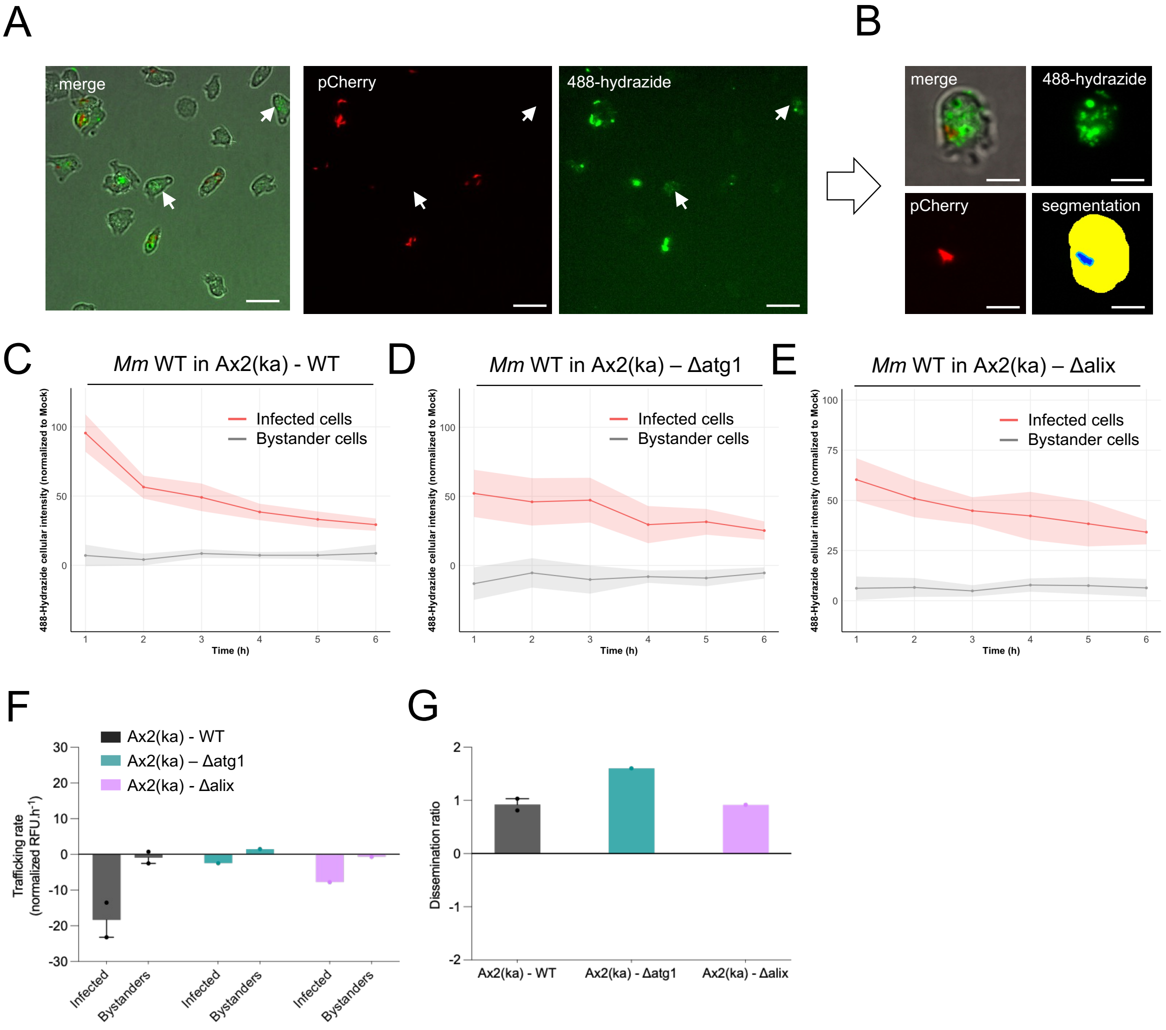


112

### Suppl. Fig. 5

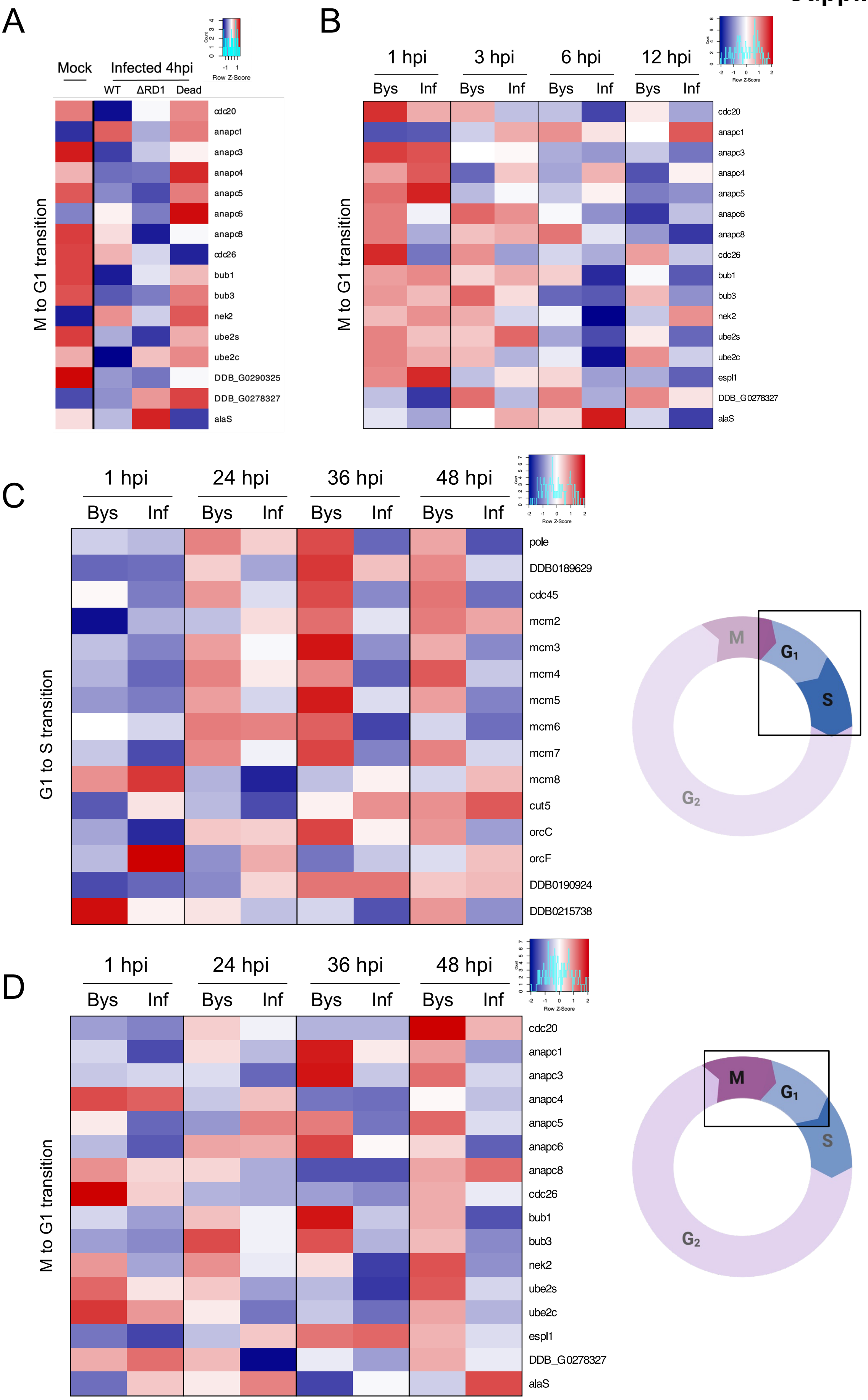


113

**Suppl. Fig. 6**

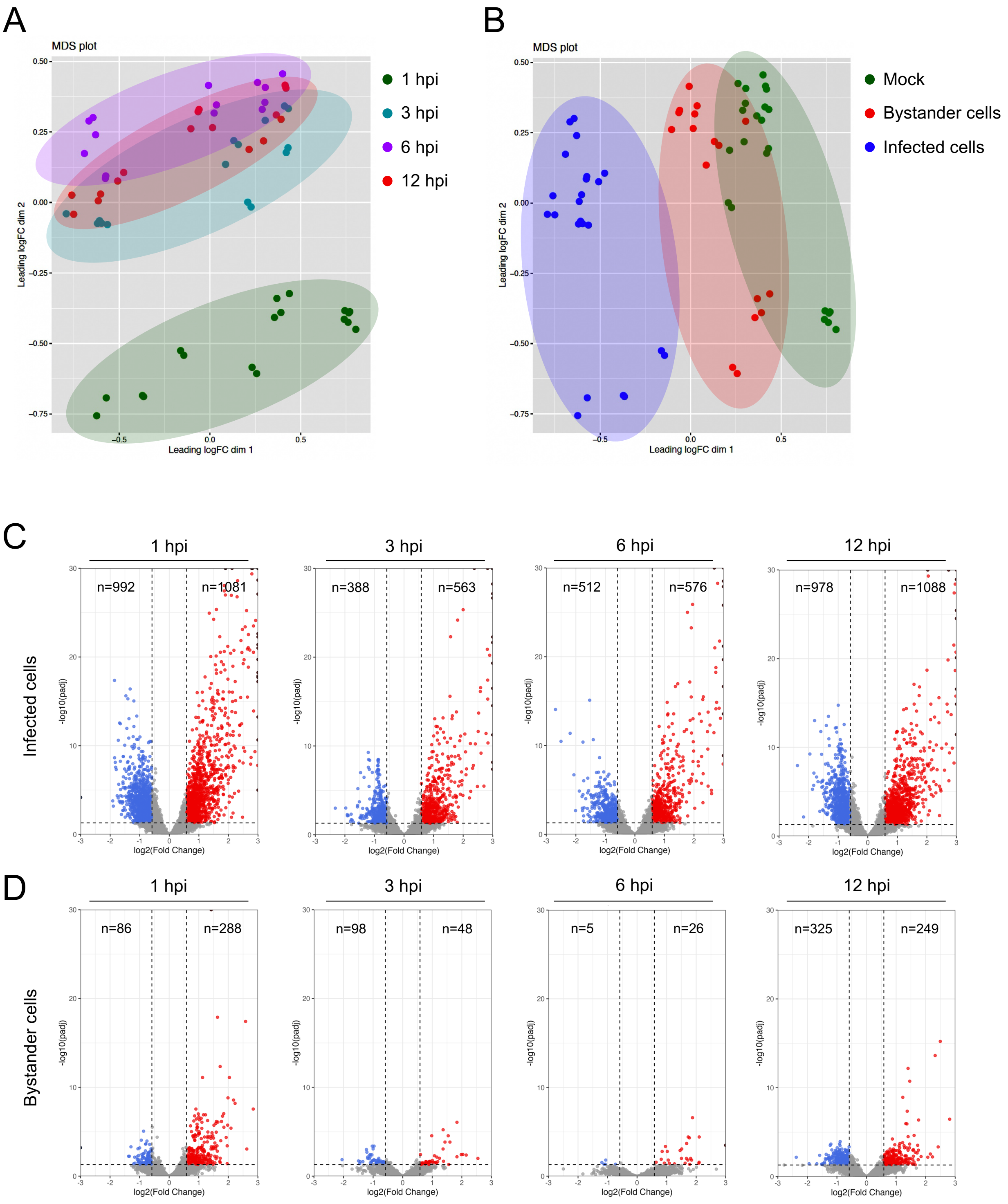


114

**Suppl. Fig. 7**

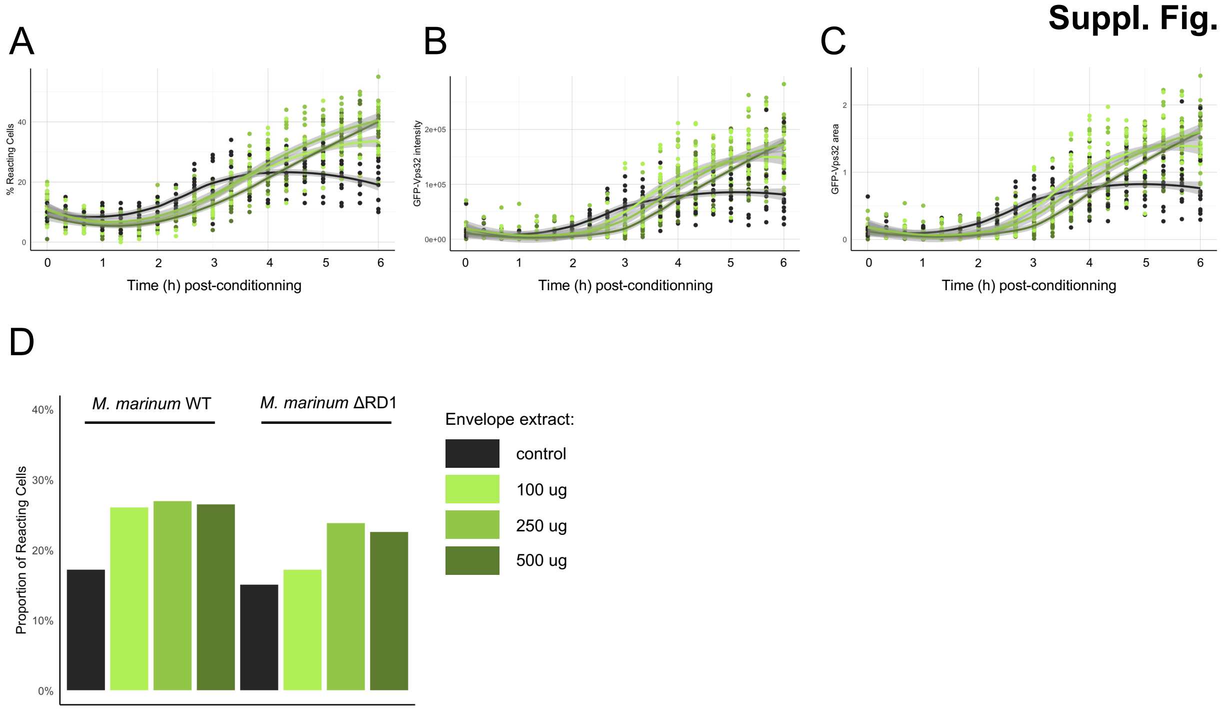
A
B
C
D
% Reacting Cells
GFP-Vps32 intensity
GFP-Vps32 area
Time (h) post-conditionning
Time (h) post-conditionning
Time (h) post-conditionning
Proportion of Reacting Cells
M. marinum WT
M. marinum ΔRD1
Envelope extract:
control
100 ug
250 ug
500 ug
